# Control of Neutrophil activation through Semaphorin 7A-Plexin C1 is essential for immune defense during pulmonary sepsis

**DOI:** 10.1101/2021.11.08.467692

**Authors:** Tiago Granja, David Köhler, Linyan Tang, Philip Burkhardt, Ka-Lin Heck-Swain, Michael Koeppen, Harry Magunia, Maximilian Bamberg, Franziska Konrad, Kristian Ngamsri, Anika Fuhr, Marius Keller, Helene A. Haeberle, Tamam Bakchoul, Alexander Zarbock, Bernhard Nieswand, Peter Rosenberger

## Abstract

Pulmonary defense mechanisms are critical for host integrity during the early phase of pneumonia and sepsis. These processes are fundamentally dependent on the activation of neutrophils during the early phase of the innate immune response. Recent work has shown that semaphorin 7A (Sema7A) holds significant impact on platelet activation, yet its role in neutrophil migration and function is not well known. We report here that Sema7A binds to neutrophil PlexinC1, increasing integrins and L-selectin on the neutrophil surface. Sema7A-induced neutrophil activation also prompted neutrophil chemotaxis in vitro and the formation of platelet-neutrophil complexes in vivo. We also observed altered adhesion and transmigration of neutrophils in *Sema7A-/-* animals in the lung. *Sema7A-/-* animals also showed altered crawling properties of neutrophils. This resulted in increased number of neutrophils in the interstitial space of *Sema7A-/-* animals but reduced numbers of neutrophils in the alveolar space during pneumonia-induced pulmonary sepsis. This was associated with significantly worse outcome of *Sema7A-/-* animals in a model of *Klebsiella pneumoniae*. Furthermore, we were able to show a correlation between serum levels of Sema7A in patients with ARDS and oxygenation levels. Thus, we show here that Sema7A has an immunomodulatory effect though which might influence patient outcome during pulmonary sepsis.

**Summary:** Sema7A controls pulmonary immune defense

## INTRODUCTION

Sepsis remains one of the leading causes of death worldwide and one of the leading causes of hospitalization (*1*). Pneumonia is frequently the source of pulmonary sepsis, and is caused by the invasion of pathogens entering the lung and eventually through the pulmonary barrier into the human circulation. In hospitalized and ventilated patients, gram-negative bacteria are frequently the cause of pneumonia, whereas gram-positive bacteria are more common in patients with community-acquired pneumonia (*2*). The treatment of pneumonia consists of fluid administration, bed rest and aggressive antibiotic therapy; however, this approach is not always successful, and the severity of the disease progresses. Therefore, understanding the mechanisms that control, influence or support the host response during pneumonia might help to improve the outcome of patients with pulmonary inflammation and infection (*3*).

Neutrophils are the forefront in the defense against pathogens that challenge the lung (*4*). Following activation, neutrophils tether and roll along the endothelial surface. This process is dependent on L-selectin expression (*5*). L-selectin cleavage from the neutrophil surface is triggered by integrin engagement and is involved in neutrophil recruitment to the lung and bacterial clearance (*6*). P-selectin expression and MAC-1 are also important during the recruitment of neutrophils to the lung and correlate with outcome in patients with ARDS (*7, 8*). Subsequently, integrins play key roles in the transendothelial migration and activation of neutrophils. The differential context of the integrins is demonstrated by the fact that blocking antibodies against the β2 integrin CD18 result in an increased number of neutrophils in the alveolar space and a reduction in pulmonary injury (*9*). β2 integrin binds to ICAM-1 on endothelial cells, which results in increased transendothelial neutrophil migration, an essential mechanism of pulmonary host defense (*10*).

Recent work has shown that the neuronal guidance protein semaphorin 7A (Sema7A) is important during the initial stages of inflammation. During the inflammatory response, Sema7A can bind to integrin α1β1 to facilitate cytokine storm (*11*). We have shown in the past that Sema7A is induced during periods of hypoxia and activates platelets through the glycoprotein Ib-IX-V (GPIb) receptor (*12, 13*). Sema7A was also reported in the past to influence pulmonary inflammation, although the exact mechanisms have not been described to date. To highlight the role of Sema7A during the pulmonary immune response, we investigated the role of Sema7A during LPS-induced and *Klebsiella pneumoniae*-induced pulmonary effects. We demonstrate that Sema7A engages the PlexinC1 receptor on neutrophils and alters the expression of key neutrophil integrins. This expression is essential for the coordinated migration of neutrophils into the alveolar space and influences the formation of platelet-neutrophil complexes, leading to significant alterations in pulmonary immune defense in a model of *Klebsiella pneumoniae*-induced pulmonary infection.

## METHODS

### Animal Ethics Statement

All animal procedures conformed to the German guidelines for the use of living specimens and were approved by the Institutional Animal Care and the Regierungspräsidium Tübingen and Würzburg.

### Human samples

Approval for the processing of human samples was given by the Ethics Committee (Institutional Review Board) of the University of Tübingen (approval number 156/2016BO1, Clinicaltrial.gov: NCT02692118). Informed consent was obtained from patients or legal guardians before samples were collected and processed for further analysis. Demographic data and laboratory values were collected from a routine patient data management system. Human samples were analyzed by flow cytometry were collected and stored for subsequent ELISA analysis.

### Mice

*Sema7A*^*-/-*^ mutant mice were generated, characterized and validated as described in Pasterkamp et al. (*14*). We generated the Sema7A floxed mouse line (*Sema7a*^*loxP/loxP*^) (*12*), which was then crossbred with the listed Cre recombinase-positive mouse lines to generate mice with the following tissue-specific gene deletions: immune cell-specific LysMCre^+^; megakaryocyte- and thrombocyte-specific PF4Cre^+^ and endothelial-specific Tie2Cre^+^. As internal experimental controls, Sema7^loxP/loxP^ Cre^-^ littermates were used. WT animals were obtained from inbreeding colonies maintained by operational researchers and animal facility staff.

### Murine lung injury model with LPS inhalation

Mice were exposed to 0.5 mg/ml LPS by inhalation for 45 min, followed by a 4-h incubation to induce acute lung injury as described previously (*15, 16*). In the experimental subsets, animals were treated with 2 µg of recombinant murine Sema7A (R&D # 1835-S3-050) or 2 µg of anti-murine Sema7A antibody (R&D # AF1835) i.v. prior LPS inhalation or 0.9% NaCl as a control treatment. Animals were littermate-matched according to sex, age, and weight, and after the appropriate incubation times, the animals were anesthetized with pentobarbital sodium (80 mg/kg BW, intraperitoneally) prior to harvesting BALF and/or blood and lung samples.

### Murine lung injury model with *Klebsiella pneumoniae* instillation

Briefly, mice were anesthetized with a 3-component fentanyl mixture applied i.p., followed by a small skin incision and exposure of the trachea. Direct tracheal instillation of 4×10^7^ *Klebsiella pneumoniae* (ATCC strain 43816) in 50 µl of PBS was performed using a 30-gauge needle to minimize tracheal damage. The incision was treated locally with carprofen to relieve pain and was then closed. Twenty-four hours after instillation, the animals were anesthetized, and the BALF, organs and blood were harvested for further analysis and histological assessments. In a similar experimental setup, the mice were not sacrificed after 24 h. PMNs in the BALF were counted and processed by flow cytometry. Additionally, agar plates (BD, Difco™ Nutrient Agar 213000) were inoculated with BALF and blood and cultured in an incubator for 24 h to grow CFUs for quantification.

### In vivo migration assay

Twenty-four hours after *Klebsiella pneumoniae* instillation, a fluorescent APC-conjugated Ly6G antibody (clone 1A8) was injected via the tail vein to label intravascular PMNs. To remove nonadherent leukocytes, saline was injected into the beating heart. PMNs from the alveolar space were examined in the BALF. The lungs were incubated with anti-CD45 PerCP-Cy5·5 (clone 30-F11) and anti-Ly6G PE/Cy7 (clone 1A8). The absolute cell counts in the BALF and lungs were determined. We differentiated between interstitial PMNs (CD45-PerCP-Cy5·5^+^; Ly6G-PE-Cy7^+^ and Ly6G- APC^-^) and intravascular PMNs (CD45-PerCP-Cy5·5^+^; Ly6G-PE-Cy7^+^ and Ly6G-APC^+^) (all antibodies from BioLegend) by flow cytometry (FACS Canto II; BD Biosciences).

### Chemotaxis Assay

Freshly isolated neutrophils were enriched with a MACS LS column system with anti-CD15 magnetic antibodies and collected in RPMI-based medium (1% Pen Strep, 1% L-glutamine, 10% FBS - PAA). Chemotaxis flow chambers (IBIDI µSlides) were coated as described by the manufacturer’s protocol with gradients of 1 µg/ml Formylmethionyl-leucyl-phenylalanine (fMLP; Sigma #F3506) and 10 µg/ml recombinant Sema7A (rhSema7A; R&D #2068-S7). Concentrated neutrophils (3×10^6^ cells/ml) were loaded into the µSlides, and microscopic images were acquired in an incubation chamber maintained at 37°C and 5% CO_2_ for 3 h. Videos were assembled from the recorded images (10-min time lapse, 20× lens, resolution of 0,32 µm/pixel, frame of 1388×1040), which were acquired on an AxioCam 305 mono (Zeiss) and an LSM 510 laser confocal microscope (Zeiss). All chemotaxis tracks were generated with the ImageJ free software tool for the analysis of time stacks (ImageJ version 1.52a, NIH-USA) according to the manufacturer’s protocol.

### Intravital microscopic analysis of cremaster microcirculation

Mouse cremaster preparation was performed as previously described (*17*). Cremaster preparation was initialized deeply anesthetizing each specimen (80 mg/kg BW pentobarbital sodium, i.p.). On a microscope with a heated table, the cremaster muscle was exposed as a flat trapeze. The prepared specimen was placed under a 20× water dipping lens (resolution of 0,32 µm/pixel) mounted on a Nikon Eclipse Ci-L microscope (Nikon, Düsseldorf; Germany) and controlled by NIS elements AR software.

After preparation, PMNs and platelets were visualized by applying 20 μl of platelet-specific FITC-labeled X488 antibody (Emfret, Eibelstadt, Germany) and 20 µl of PE-labeled anti-Ly6G (BioLegend #127608) with a catheter. Images were acquired by a Hamamatsu Orca Flash 4.0 camera mounted on a dual emission image splitter (optoSplit II, Cairn Research; UK) at a rate of 16 frames/sec (on videos of 10 sec total) and a resolution of 2048 ×1024 pixels. The mice were treated i.v. with LPS (Sigma; 0.4 μg/g BW). All analyses and data collection were performed using NIS elements Advanced Research software (Nikon, Düsseldorf, Germany).

### Lung intravital microscopy

Anesthesia was performed with ketamine/xylazine (100 mg/kg/16 mg/kg), and an antibody cocktail composed of 7 µg of anti-CD31-A647, 5 µg of Ly6G-A488, and 5 µg of GPIX-A546 was administered i.v. in a bolus of 100 µl. For correct microscopy of the lung, the mouse was intubated by tracheotomy, and breathing was normalized for 5 min after the instillation of 5 µg/g BW LPS (O26:B6). For murine lung microscopy, a Leica Stellaris 8 resonance microscope with a 25× objective and a resolution of 232.72 µm^2^ was used. Acquisition was performed for 10 min for a total time of 30 min with and without recombinant Sema7A (recSema7A; R&D #AF1835). Videos were analyzed with Leica software.

### Flow cytometry

All measurements were performed with a FACSCanto II flow cytometer (BD, Heidelberg, Germany), and all instrument calibration, performance and plotting of data were executed with BD FACSDiva software (version 6). Detailed data analysis of the acquired samples was performed using FlowJo software (version 10x and above, Ashland, Oregon, USA).

### Staining of murine platelet-neutrophil complexes

Whole blood was gently collected from the left ventricle of the heart with a 25G needle in a 1:10 citrate-coated syringe and incubated at 37°C in warmed tubes with an antibody cocktail. After with the addition of BD lysis buffer (BD, #349202), the cells were incubated with an antibody cocktail of anti-CD42b FITC (Emfret #M040-1), anti-CD66b PE/Cy7 (BioLegend #108416), anti-LFA1-PE (BioLegend #141006), anti-CD11b BV421 (BioLegend #10236), anti-CD62P-APC (BD #563674), anti-CD62L PerCP/Cy5.5 (BioLegend #104431) and rat anti-mouse activated GPIIb/IIIa PE (Emfret, clone JON/A #M023-3) prior to acquisition by flow cytometry.

### Human ICAM-1 binding assay

Human neutrophils were freshly isolated with a MACS LS column system (enriched with magnetic anti-CD15 antibodies) as described previously, collected in HBSS^+/+^ and used as controls (3×10^5^ cells/ml) or were incubated with 10 ng/ml TNFα (PeproTech #AF-300-01A), 10 µg/ml recSema7A Fc (R&D 2068-S7), or CD11b antibodies (Invitrogen #14-0112-82). Under these different conditions, cells were incubated with soluble recombinant human ICAM-1/CD54 Fc chimeric protein (100 ng/ml) (R&D, # 720-IC-050) for 10 min at 37°C. The experiment was stopped with 500 µl of 1× BD cell fix solution, and after being washed, the cells were labeled with anti-CD66b-PE/Cy7 and anti-human CD54-AF647 antibodies. Experimental cell viability was assessed by the exclusion of DRAQ7 (Biolegend # 424001), and apoptotic cells were examined with anti-Annexin V (BD # 556547).

### Regulation of adhesion receptors

Human neutrophils were isolated on a MACS LS column system (Miltenyi Biotec #130-042-401), enriched with magnetic anti-CD15 antibodies (Miltenyi Biotec #130-046-601) and collected in HBSS^+/+^ buffer. To observe the shedding of SEMA7A from the surface of PMNs, in one set of experiments, PMNs (2.5×10^6^ cells/ml) were used as a control or stimulated with 10 ng/ml TNFα (PeproTech #AF-300-01A) for 15 min at 37°C. The reaction was stopped with 1× BD Cytofix/Cytoperm solution (BD, Kit #554722), and the cells were washed and incubated with anti-human CD66b FITC (Biolegend #305104) and anti-human Semaphorin 7A-AF647 (R&D #FAB20681R) for 30 min at RT.

To examine the regulation of PMN adhesion surface receptors, we tested the effect of 10 µg/ml human recombinant Sema7A Fc (R&D #2068-S7) for 15 min at 37°C. The cells were fixed with 1× BD Cytofix/Cytoperm solution (BD, Kit # 554722), washed and incubated with anti-human CD66b PE/Cy7 (eBioscience #25-0666-42), CD11b PE (Beckman Coulter # PN-IM2581U), anti-human LFA1 (CD11a) FITC (Biolegend 363404) and anti-human PSGL1 BV421 (BD # 563961) for 30 min at RT.

### Ex vivo capillary flow chamber assay

Neutrophil adhesion under flow was measured by perfusing human whole blood on rectangular inner profile boro tubing glass capillaries (VitroCom, # 427411) coated with 3.0 µg/ml recombinant human E-selectin/CD62E Fc chimeric protein (R&D # 724-ES) or on a combination of E-selectin and 3.5 µg/ml recombinant human ICAM-1 (CD54 Fc chimera protein; R&D # 720-IC). Stabilized capillaries were imaged on a Leitz microscope with a ×lens (NA=0.32) equipped with a black-and-white Hamamatsu Orca R^2^ camera (Hamamatsu, Herrsching-Germany), and 10-sec videos were recorded with Nikon NIS Elements Advanced Research software (Nikon, Düsseldorf, Germany). All cells were tracked manually with Nikon NIS Elements Advanced Research software and the free ImageJ software tool for cell tracking analysis (ImageJ version 1.52a, NIH-USA).

### Immunofluorescent staining of purified murine PMNs

After purification, PMNs were washed and fixed on silanized slides on a hot plate for 10 min at 60°C. After being blocked with donkey serum (1:20 dilution) for 30 min, the slides were incubated with an antibody cocktail composed of anti-mouse CD11b (1:50 dilution, rat monoclonal IgG2b, eBioscience, #14-0112-86) and anti-mouse semaphorin 7A (1:20 dilution, polyclonal goat IgG, R&D #AF1835) combined with anti-mouse EGFR (1:50 dilution, rabbit IgG, Cell Signaling Catalog # 4267), anti-mouse PlexinC1 (1:20 dilution, polyclonal sheep IgG; R&D # AF5375) or anti-mouse integrin β1 (1:100 dilution, CD29; rabbit mAb Cell Signaling #34971). The slides were counterstained with DAPI and observed under a Zeiss LSM 510 confocal microscope operated with ZEN software Black edition (Zeiss).

### Immunofluorescent staining of murine lung tissue

After LPS inhalation and incubation for 4 h, murine lung tissues were fixed in 4% paraformaldehyde and embedded in paraffin blocks prior to prepared as 3 µm slices. After being blocked with goat serum, rabbit anti-mouse Sema7A primary antibodies (Abcam, ab31449) were added and incubated, followed by goat anti-rabbit Alexa 488 secondary antibody staining. Nuclear counterstaining was performed with DAPI. Lung sections were observed on a Zeiss LSM510 with an 63× oil magnification lens, and images were taken with an Axiocam (Zeiss) and processed by ZEN software (Zeiss).

### Immunofluorescent staining of human PMNs

PMNs were incubated with fMLP or NaCl (control) for 15 min, fixed with 4% PFA for 5 min at RT and mounted on a slide at 60°C for 10 min. For IF staining, we used the primary antibodies anti-CD45 (rabbit anti-human; Santa Cruz #sc-25590) and anti-Sema7A (mouse anti-human; Santa Cruz #sc-374432). PMNs were examined on a Stellaris 8 (Leica Microsystems) with an oil 63× oil magnification lens using LAS X software 3.7.3.23245 (Leica Microsystems).

### ELISA

All enzyme-linked immunosorbent assay (ELISA) kits were from R&D and used the DuoSet principle. The absorbance of the developed color was measured at 450 nm in a plate reader (Tecan, Männedorf, Switzerland).

### Respiratory burst assay

The respiratory burst assay was based on the conversion of rhodamine 123 (5 mg/ml, a 1:1000 dilution) by reactive species produced by neutrophils in whole blood. Samples were collected from the antecubital veins of healthy donors and incubated for 15 min at 37°C with 10 ng/ml recombinant human TNFα (Promokine #C-63721), 2 µg/ml recombinant Sema7A (R&D #2068-S7) or 2 µg/ml anti-PlexinC1 antibody (R&D #3887). The reaction was stopped, and each sample was measured by flow cytometry at 530 nm on a FACSCanto II.

### Phagocytosis assay

Phagocytosis was analyzed in human whole blood that was treated for 15 min with 10 µg/ml recombinant human Sema7A (R&D 2068-S7), 10 ng/ml TNF (PeproTech AF-300-01A), or PlexinC1 blocking antibodies (R&D, # MAB3887) at 37°C or at 4°C as a negative control. Whole blood samples were incubated with pHrodo Green *E. coli* BioParticles Conjugate (Thermo Fisher # P35366) (Ex/Em 509/533), and after incubation, the blood cells were lysed with 1× BD lysis buffer (BD, 349202) prewarmed to 37°C. After being centrifuged at 500 g for 5 min, the cells were resuspended in RPMI containing Molecular Probes™ Vybrant™ DyeCycle™ stain (Thermo Fisher # V10309), incubated for 15 min on ice and fixed with 1× BD cell fix solution (BD #340181). After being washed with ice-cold HBSS^-^, the cells were incubated with BV421-conjugated anti-human CD11b (Biolegend # 301324) and APC-conjugated anti-human PMN Ly6G/6C antibodies (Biolegend # 108412) in PBS containing 1% BSA and 0.05% sodium azide. The stained samples were analyzed in a FACSCanto II flow cytometer (BD, Heidelberg, Germany).

### Proteomics analysis

Proteomics analysis of proteins and phosphorylation was performed by Sciomics scioDiscover antibody microarrays. Proteins were extracted from stimulated and unstimulated neutrophils with scioExtract buffer (Sciomics), and the bulk protein concentration was determined by BCA assays. Samples were labeled with scioDye2, and the reaction was stopped after 2 h with excess dye and washed with buffered PBS. The arrays were blocked with scioBlock (Sciomics) on a Hybstation 4800 (Tecan, Austria). Slide scanning was conducted using a Powerscanner (Tecan, Austria), and spot segmentation was performed with GenePix Pro 6.0 (Molecular Devices, Union City, CA, USA). The acquired raw data were analyzed by Sciomics using the linear models for microarray data (LIMMA) package of R-Bioconductor after uploading the median signal intensities. For normalization, cyclic loess normalization was applied to all acquired data. To analyze the samples, a one-factorial linear model was fitted with LIMMA, resulting in a two-sided t-test or F-test based on moderated statistics.

### Immunohistochemistry

For immunohistochemical staining of platelet-neutrophil complexes in experimental imbedded lung tissue sections, we used the Vectastain ABC Kit (Linaris, Wertheim, Germany) to analyze the fixed lungs 24 h after Klebsiella instillation.

### H&E staining

Cryosections of lung tissues were placed in PBS at RT for 10 min, fixed in ice-cold methanol and washed 2 times in Aqua dest. Staining was performed as described previously (*12*).

### Statistical analysis

The data are presented as bar graphs using the mean ± SD. Statistical analysis was performed using Student’s *t*-tests to compare two groups; for comparison of multiple groups, we performed one-way analyses of variance followed by Dunnett’s tests. For all performed comparisons, *p*-values are displayed, and *p*-values < 0.05 were considered statistically significant (displayed as *p* < 0.05 (^*^); *p* < 0.01 (^**^) and *p* < 0.001 (^***^)).

## RESULTS

### Sema7A is expressed at the pulmonary immune interface and affects neutrophil migration

We have previously shown that Sema7A is induced during inflammatory hypoxia. Whether Sema7A is important for pulmonary defense during sterile or nonsterile inflammation at the pulmonary immune interface is not known. Therefore, we used *Sema7A*^*-/-*^ mice and littermate controls to establish a model of *Klebsiella pneumoniae*-induced pneumonia and found significantly altered histological images with a stronger changes in the interstitial and alveolar structures in the *Sema7A*^*-/-*^ mice compared to those of the littermate controls (**Figure 1A**). We then stained Sema7A in pulmonary tissue and on neutrophils, since these are the first line of defense at the pulmonary alveolar-capillary barrier during an external assault, and found that Sema7A was clearly expressed on neutrophils and the vascular barrier within the lung (**Figure 1B and C**). Neutrophil migration to the site of infection is another important mechanism in the defense of the pulmonary surface; therefore, we exposed *Sema7A*^*-/-*^ mice and littermate controls to i.v. LPS injection and performed intravital microscopic analysis of a cremaster model staining neutrophils and platelets. We found significantly increased cell speed, reduced numbers of stationary cells and reduced numbers of transmigrated neutrophils in *Sema7A*^*-/-*^ animals compared to the controls (**Figure 1D and E**). We then repeated this analysis in the pulmonary circulation following LPS inhalation by injecting recSema7A to determine whether we could visualize the effect of Sema7A within the lung. We found an increase in the area that was covered with neutrophils in the pulmonary circulation and an increase in the number of platelet-neutrophil complexes in recSema7A-injected animals (**Figure 1F and G**).

**Figure 1:**
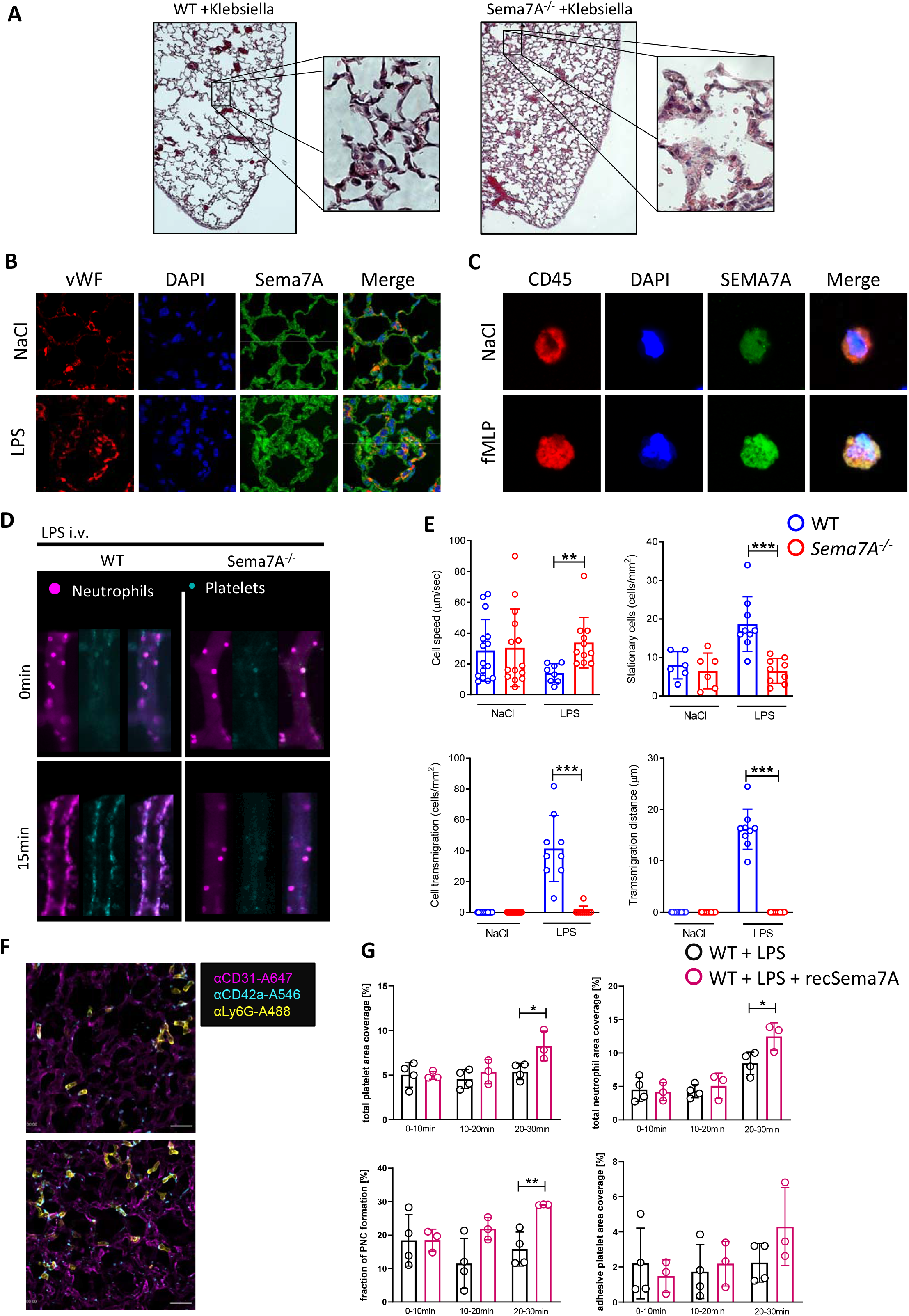
Semaphorin 7A is required for neutrophil adhesion and migration during inflammation. **A)** Histological cross-sections and magnifications of lung tissue from WT and *Sema7A*^*-/-*^ mice 24 h after the instillation of 4×10^7^ *Klebsiella pneumoniae* cells. **B)** Immunofluorescence (IF) staining of Sema7A (green) and vWF (red) in endothelial cells and nuclear staining with DAPI (blue). **C)** IF staining showing Sema7A expression (green) on the surface of Ly6G-marked PMNs (red) collected from the peripheral blood of WT mice treated with LPS inhalation and 4 h of incubation. **D)** Representative images of platelets (cyan) and neutrophils (magenta) captured by intravital microscopy videos of murine cremaster tissues from WT and *Sema7A*^*-/-*^ specimens after 15 min of incubation with 1 mg/ml LPS. **E)** PMN speed, stationary PMNs at the vessel wall, transmigrated PMNs and the distance after transmigration were evaluated in multiple video sequences and collected from independently performed triplicate experiments. **F)** Total neutrophil area coverage, total and adhesive platelet area coverage and the fractions of PNC formed in the lung, as determined via intravital confocal microscopic analysis of the lung in WT mice instilled with 5 µg/g BW LPS, with or without additional treatment with recombinant Sema7A. In **E)** and **F)**, only relevant group comparisons were performed by unpaired two-tailed Student’s t-tests (the data are the mean ± SD). ^*^p < 0.05, ^**^p < 0.01 and ^***^p < 0.001 as indicated.

### Sema7A binds to PlexinC1 on neutrophils

Next, we identified which receptor Sema7A binds to during inflammatory stimulation to better understand the underlying principles and mechanisms of this process. We isolated neutrophils from WT animals following NaCl and LPS inhalation and examined the localization of Sema7A during inflammatory stimulation of neutrophils. We stained for the Sema7A receptors integrin β1 (CD29), PlexinC1 and the EGF receptor, since all of these factors have been reported to be potential target receptors for Sema7A. We found a robust Sema7A signal on neutrophils after LPS treatment, as indicated by staining for PlexinC1 and Sema7A (**Figure 2C**), whereas no co-expression of Sema7A with CD29 or EGFR was observed (**Figure 2A and C**). To further validate this finding, we injected recSema7A into *Sema7A*^*-/-*^ mice and isolated neutrophils. Following injection of recSema7A in *Sema7A*^*-/-*^ animals, we found robust colocalization of Sema7A with the PlexinC1 receptor on these cells, showing the direct binding of recSema7A to PLXNC1 (**Figure 2D**).

**Figure 2:**
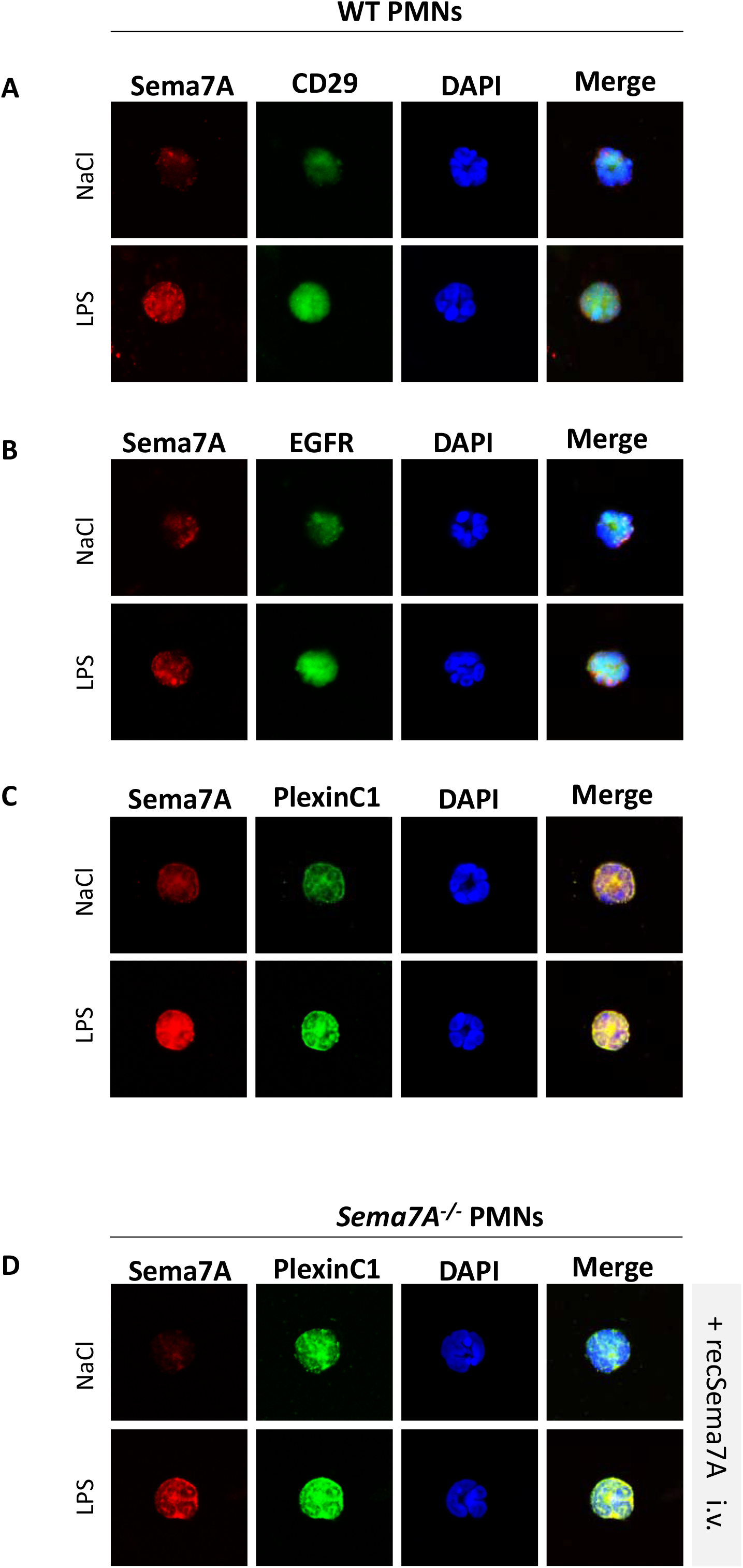
Sema7A binds to the neutrophil PlexinC1 during LPS inhalation. Stained neutrophils isolated from saline (NaCl)- or LPS-treated WT and *Sema7A*^*-/-*^ mice 4 h after incubation. **A)** Expression of Sema7A (red) and CD29 (green) on PMNs harvested from WT mice treated with NaCl or LPS. **B)** Expression of Sema7A (red) and EGFR (green) on PMNs harvested from WT mice treated with NaCl or LPS. No protein colocalization was visible in the merged pictures in either condition. **C)** Expression of Sema7A (red) and PlexinC1 (green) on PMNs harvested from WT mice treated with NaCl or LPS. Sema7A expression is highly increased in LPS-treated mice, and the merged pictures show a strong interaction between Sema7A and PlexinC1. **D)** Surface PMN expression of Sema7A (red) and PlexinC1 (green) in *Sema7A*^*-/-*^ mice after the injection of exogenous recombinant Sema7A. Strong binding of exogenous Sema7A to *Sema7A*^*-/-*^ PMNs was observed. The merged pictures show an intense correlation with PlexinC1. Multiple acquisitions of stained cells were analyzed from independently performed triplicate experiments.

### Sema7A induces neutrophil migration through PlexinC1

Chemotaxis is an essential step by which neutrophils migrate to the site of infection and/or inflammation. This step is crucial in the host defense against pathogens. Therefore, we investigated whether Sema7A influenced neutrophil migration during a chemotactic stimulus. We used an in vitro cell migration assay with neutrophils exposed to fMLP or Sema7A or neutrophils preincubated with anti-PLXNC1 antibody before Sema7A exposure. Treatment with fMLP resulted in the attraction of neutrophils towards the highest concentration, with profound migratory velocity and distance (**Figure 3 A-E**). In contrast, recSEMA7A resulted in the repulsion of neutrophils, with similar velocities and distances as the effect of fMLP. This effect could be reduced when neutrophils were preincubated with anti-PLEXINC1 antibodies before the experiment (**Figure 3A-E)**. This finding showed that neutrophil migration was dependent on Sema7A and that PlexinC1 could reverse the effect of Sema7A. Next, we examined whether Sema7A affected the binding of fibrinogen to MAC-1 to determine Sema7A-mediated changes in the affinity of this receptor. We found that Sema7A highly affected the binding of fibrinogen to MAC-1, suggesting the significant influence of Sema7A on integrin activation (**Figure 3F**). In addition, we also performed an ICAM-1 binding assay and found that Sema7A significantly influenced the binding of ICAM-1 to neutrophils stimulated with Sema7A. This finding suggests that Sema7A influences the activity of key integrins involved in neutrophil migration (**Figure 3G**) and that Sema7A has a significant impact on these functions. In a final step, we highlighted the role of Sema7A in a flow chamber experiment. This result showed that recSEMA7A significantly influenced rolling velocity by LFA-1 activation which results in binding to an E-selectin- and ICAM-1-covered surface (**Figure 3H**).

**Figure 3:**
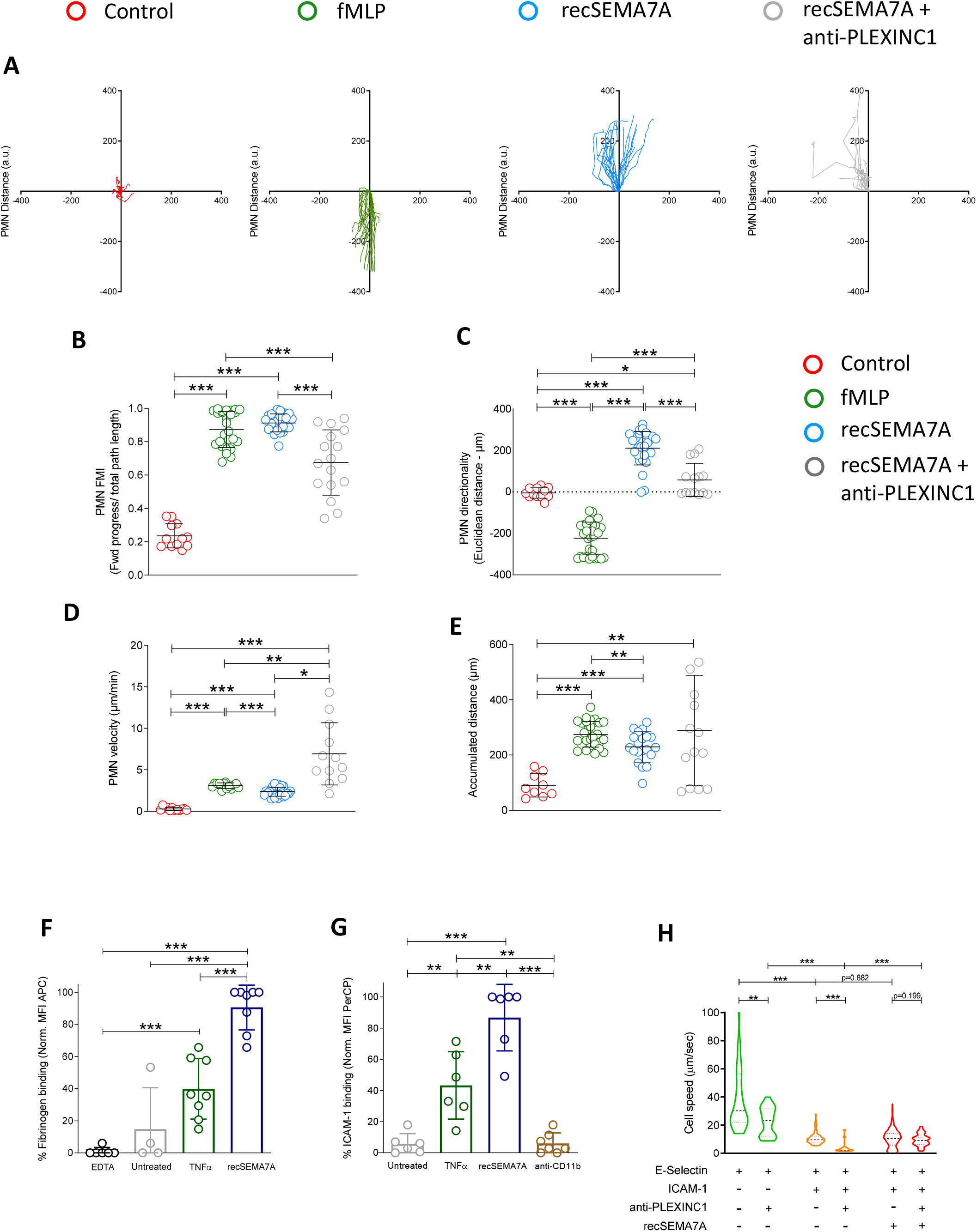
Neutrophil chemotaxis is influenced by SEMA7A through PlexinC1. Human PMNs were subjected to different stimuli in bidirectional chemotactic chambers. Acquired time lapse videos over a 3 h period were analyzed. **A)** Representative plots of PMN chemotactic tracks towards NaCl (control; red), fMLP (green), recombinant human SEMA7A (recSema7A; blue) or recSema7A together with antibodies against human PlexinC1 (anti-PLEXINC1; gray). **B, C, D and E)** Comparison of the chemotaxis parameters FMI (forward migration index), Euclidean distance under the aspect of the direction, PMN velocity and accumulated PMN distance. **F)** PMN binding affinity was indicated by APC-labeled fibrinogen on the surface of Ly6G-positive PMNs, as analyzed by FACS. The EDTA group was the internal negative control to measure the baseline autofluorescence, the untreated group was the fibrinogen negative control, TNF-α was used as a potent PMN stimulator, and treatment of PMNs with recombinant SEMA7A prior to APC-labeled fibrinogen represented the fibrinogen binding target group of interest. The fibrinogen-APC MFI was normalized and is displayed as a percentage. **G)** PMN binding affinity was indicated by PerCP-labeled ICAM-1 on the surface of Ly6G-positive PMNs by FACS. The untreated group was the ICAM-1 negative control, TNF-α was used as a potent PMN stimulator, and treatment of PMNs with recombinant SEMA7A prior to PerCP-labeled ICAM-1 represented the ICAM-1 binding target group of interest. CD11b antibody treatment was used as a control for the inactivation of ICAM-1 binding. The ICAM-1 PerCP MFI was normalized and is displayed as a percentage. **H)** Recorded neutrophil speed in µm/sec in glass flow chambers coated and blocked with casein prior to whole blood exposure to different coatings. Blocking PLEXINC1 yielded a velocity similar to that of SEMA7A coating alone. Multiple cells were analyzed from independently performed experiments in triplicate. In (B - H), all group comparisons were performed by unpaired two-tailed Student’s t-tests (the data are the mean ± SD). In (H), the p values of relevant nonsignificant comparisons are displayed. ^*^p < 0.05, ^**^p < 0.01 and ^***^p < 0.001 as indicated.

### Neutrophil proteomics analysis confirms Sema7A-induced integrin activation

To confirm the functional data and gain a better understanding of exactly how neutrophils react to Sema7A, we decided to stimulate neutrophils directly with recSEMA7A for 15 min. We analyzed surface proteins and intracellular phosphoproteins to determine which pathways were involved in the observed results. For comparison, we also exposed neutrophils to TNF-α (100 pg/ml), since TNF-α is a cardinal cytokine for neutrophil attraction. We found a significant effect of Sema7A on the expression of PSGL-1 and ICAM-1 (**Figure 4A**). This was the opposite of the effect of TNF-α and resulted in an increase in L-selectin and an increase in ICAM-1 on the surface of neutrophils. Moreover, we found a reduction in L-selectin expression, likely induced through the shedding of L-selectin. Phosphoproteomics analysis showed that the potential pathways involved in this effect were the MAP-kinase and PTEN pathways (**Figure 4B**). These pathways were previously described to be involved in the control of integrin activation on the surface of neutrophils. To confirm the obtained results, we examined some of the proteins on neutrophils through flow cytometry and found that the expression of CD11b was increased following recSEMA7A stimulation. In addition, we were able to confirm that PSGL-1 and L-selectin were reduced (**Figure 4C-F**).

**Figure 4:**
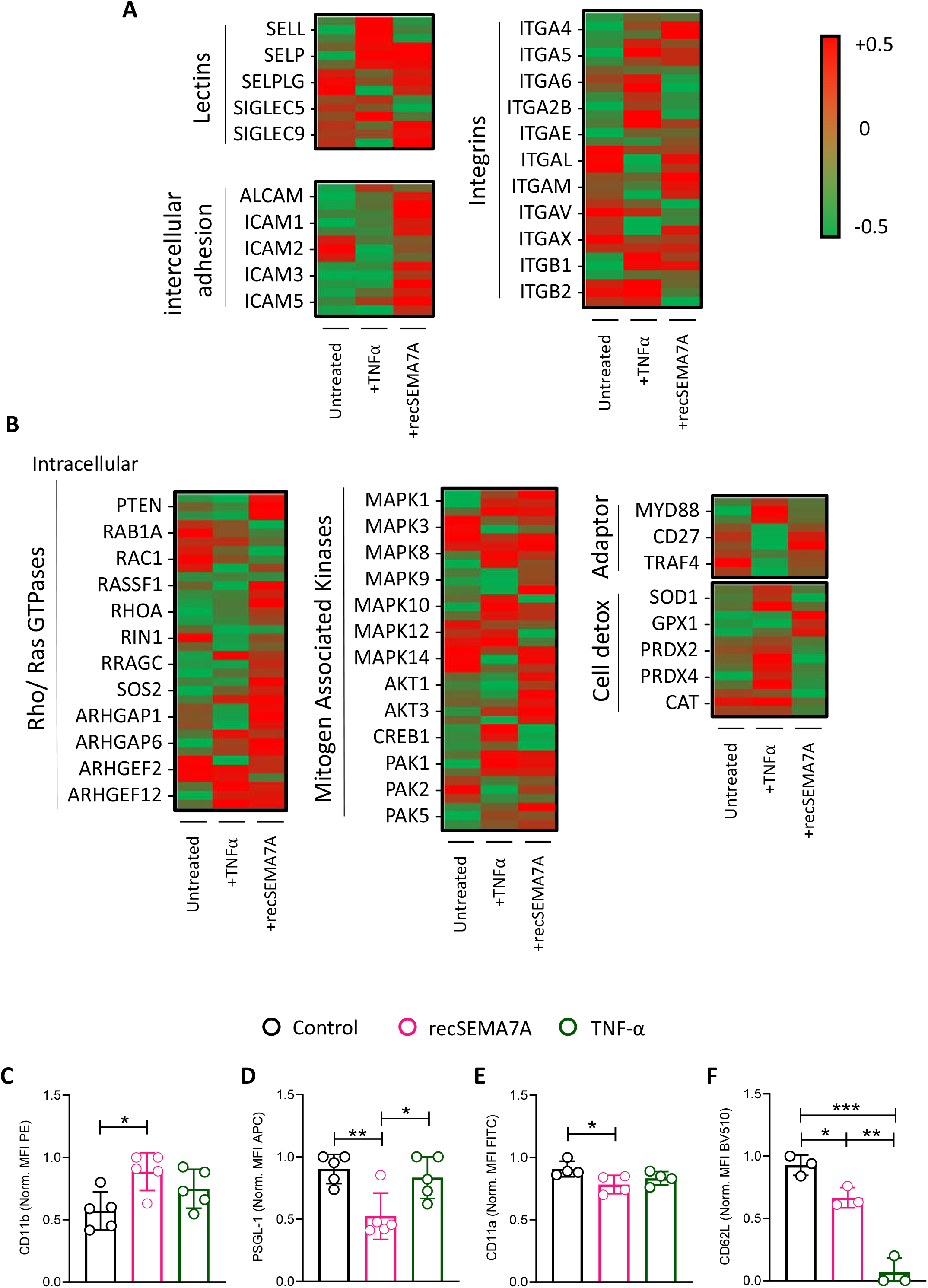
Essential neutrophil integrins are influenced by SEMA7A. Human PMNs were incubated with NaCl, 10 ng/ml TNFα or 2 µg/ml recSEMA7A for 15 min prior to snap freezing and protein extraction for proteomics analysis. Proteins were defined as differential when |logFC| > 0.5 and an adjusted p value < 0.05. The acquired raw data were analyzed by Sciomics using the linear models for microarray data (LIMMA) package of R-Bioconductor after uploading the median signal intensities. For normalization, cyclic loess normalization was applied to all acquired data. To analyze the samples, a one-factorial linear model was fitted with LIMMA, resulting in a two-sided t-test or F-test based on moderated statistics. All presented p values were adjusted for multiple analyses by controlling the false discovery rate according to Benjamini and Hochberg. Proteins were defined as differential when |logFC| > 0.5 and an adjusted p value < 0.05 from triplicate experiments. **A)** Expression of neutrophil surface lectin proteins and membrane integrin proteins from harvested samples. **B)** Expression of intracellular neutrophil Rho/Ras GTPases, mitogen-associated kinases, adaptor proteins and detoxifying enzymes. **C)** PMN surface expression of CD11b after 15 min of incubation with recSEMA7A or TNFα. Measurement was performed by FACS, and the MFI (PE) was normalized to the highest measured value. **D)** PMN surface expression of PSGL-1 after 15 min of incubation with recSEMA7A or TNFα. Measurement was performed by FACS, and the MFI (APC) was normalized to the highest measured value. **E)** PMN surface expression of CD11a after 15 min of incubation with recSEMA7A or TNFα. Measurement was performed by FACS, and the MFI (FITC) was normalized to the highest measured value. **F)** PMN surface expression of CD62L after 15 min of incubation with recSEMA7A or TNFα. Measurement was performed by FACS, and the MFI (BV510) was normalized to the highest measured value. Multiple cells were analyzed from independently performed experiments in triplicate. Group comparisons were performed by unpaired two-tailed Student’s t-tests (the data are the mean ± SD). ^*^p < 0.05, ^**^p < 0.01 and ^***^p < 0.001 as indicated.

### Neutrophil- and platelet-derived Sema7A expression alters leukocyte migration during inflammation

In the next step, we identified the source of Sema7A observed in the previous results. We have previously shown that RBCs derived from myocardial reperfusion injury are important during sterile reperfusion injury. We did not find an involvement of RBC-derived Sema7A in neutrophil migration by intravital microscopy. Next, we used *LysMCre*^*+*^*Sema7A*^*loxP/loxP*^ animals, which lack Sema7A expression on neutrophils. These mice showed significantly reduced numbers of adherent and transmigrated cells and significant differences in cell speed compared to those of littermate controls (**Figure 5A-B**). We found similar results when we examined *PF4Cre*^*+*^*Sema7A*^*loxP/loxP*^ mice in this model. These animals also showed reduced transmigration and adhesion properties in the microcirculation following LPS challenge (**Figure 5C-D)**. The possibility remained that endothelial-expressed Sema7A could bind to neutrophils and cause the observed results. When examining endothelial expression of Sema7A in *Tie2Cre*^*+*^*Sema7A*^*loxP/loxP*^ animals, to our surprise, we found no contribution of endothelial Sema7A to neutrophil attachment or transmigration (**Figure 5E-F**).

**Figure 5:**
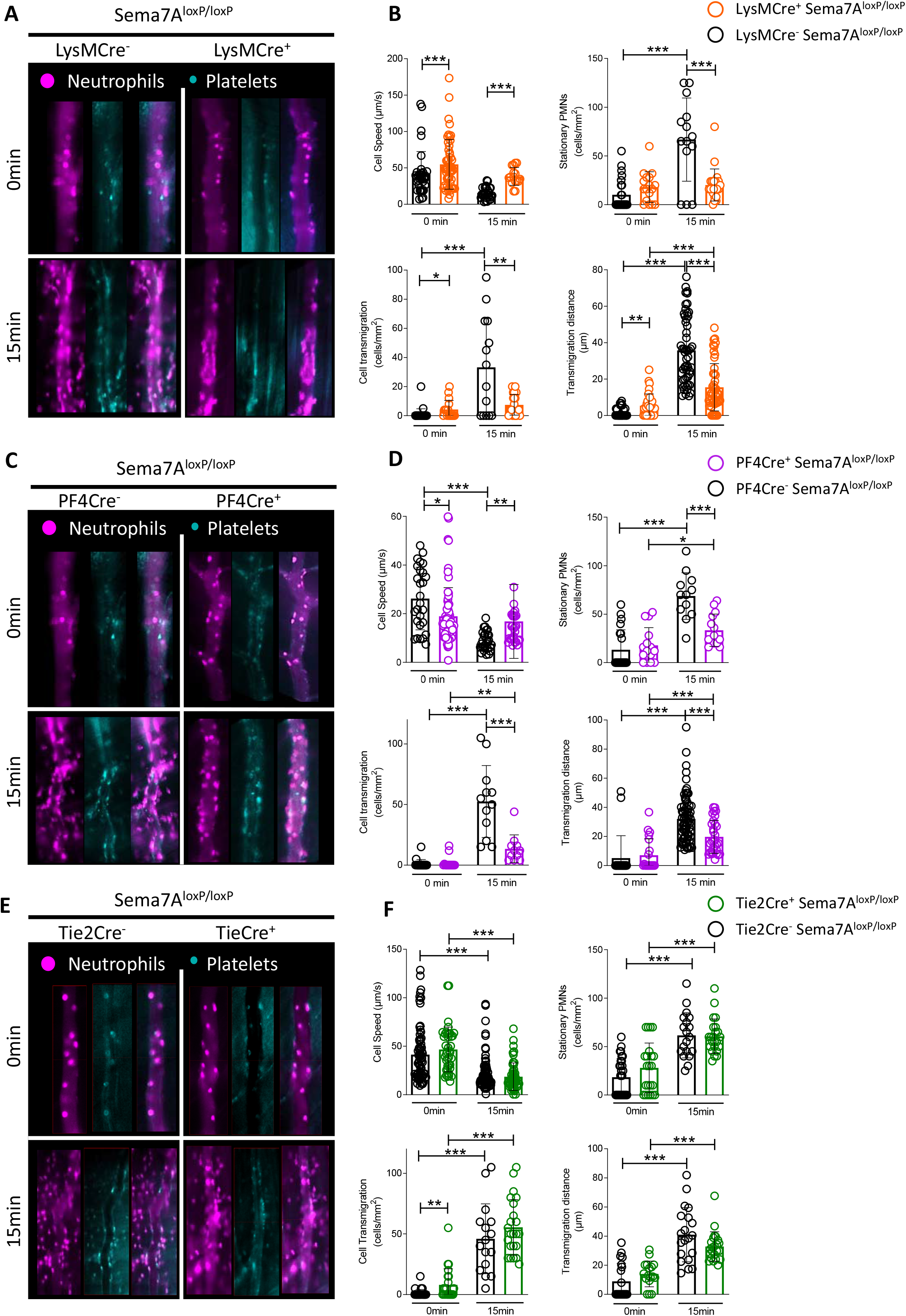
Tissue-specific expression of Sema7A controls neutrophil migration in response to inflammation. Intravital microscopic analysis of murine cremaster tissue after i.v. LPS stimulation shows the role of Sema7A expression in different cells during inflammation. **A)** Representative images of the microvasculature of *LysMCre*^*+*^*Sema7A*^*loxP/loxP*^ mice and littermate controls exposed to LPS for 15 min compared to the baseline control (0 min). **B)** Cell speed, transmigration, transmigration distance and stationary PMNs in *LysMCre*^*+*^*Sema7A*^*loxP/loxP*^ and littermate controls were analyzed by intravital microscopy after exposure to LPS for 15 min and compared to the baseline control (0 min). **C)** Representative videos of the microvasculature of *PF4Cre*^*+*^*Sema7A*^*loxP/loxP*^ mice and littermate controls exposed to LPS for 15 min compared to the baseline control (0 min). **D)** Cell speed, transmigration, transmigration distance and stationary PMNs of *PF4Cre*^*+*^*Sema7A*^*loxP/loxP*^ mice and littermate controls was analyzed by intravital microscopy after exposure to LPS for 15 min and compared to the baseline control (0 min). **E)** Representative videos of the microvasculature of *Tie2Cre*^*+*^*Sema7A*^*loxP/loxP*^ mice and littermate controls exposed to LPS for 15 min and compared to the baseline control (0 min). **F)** Cell speed, transmigration, transmigration distance and stationary PMNs in *Tie2Cre*^*+*^*Sema7A*^*loxP/loxP*^ and littermate controls was analyzed by intravital microscopy after exposure to LPS for 15 min and compared to the baseline control (0 min). Triplicate experiments were performed, and multiple cells were tracked for 15 to 20 min after LPS incubation over periods of 10 sec at 90 fps. From the acquired videos, cells were tracked manually, and relevant group comparisons were performed by unpaired two-tailed Student’s t-tests (the data are the mean ± SD). ^*^p < 0.05, ^**^p < 0.01 and ^***^p < 0.001 as indicated.

### PNC formation is significantly influenced by Sema7A

Platelet-neutrophil complexes are significant effectors of the immune response during the early phase of inflammation. We have shown that Sema7A affects PNC formation during myocardial IR injury and tissue hypoxia. However, the pulmonary barrier has to be defended against attacks by external pathogens, which try to invade through the pulmonary barrier. Since we have shown that Sema7A promotes integrin activation on neutrophils, we next performed flow cytometric analysis of WT and *Sema7A*^*-/-*^ animals 5 min after LPS inhalation. We found reduced numbers of PNCs in the blood of *Sema7A*^*-/-*^ animals without reduced expression of platelet or neutrophil activation markers (**Figure 6A-B**). We next injected recSema7A into *Sema7A*^*-/-*^ animals to determine whether we could restore this effect in animals with Sema7A gene deletion. We found that following injection of 1 µg/mouse recSema7A, the formation of PNCs in the blood was restored in *Sema7A*^*-/-*^ animals (**Figure 6C-D**). To test whether this effect could be counteracted, we injected anti-Sema7A antibodies and IgG controls at the start of LPS inhalation. Anti-Sema7A antibodies clearly reduced the number of PNCs in the vasculature of experimental animals compared to IgG-treated animals (**Figure 6E-F**). These data clearly demonstrate the PNC-forming properties of Sema7A, which are largely mediated by integrin activation on neutrophils through Sema7A.

**Figure 6:**
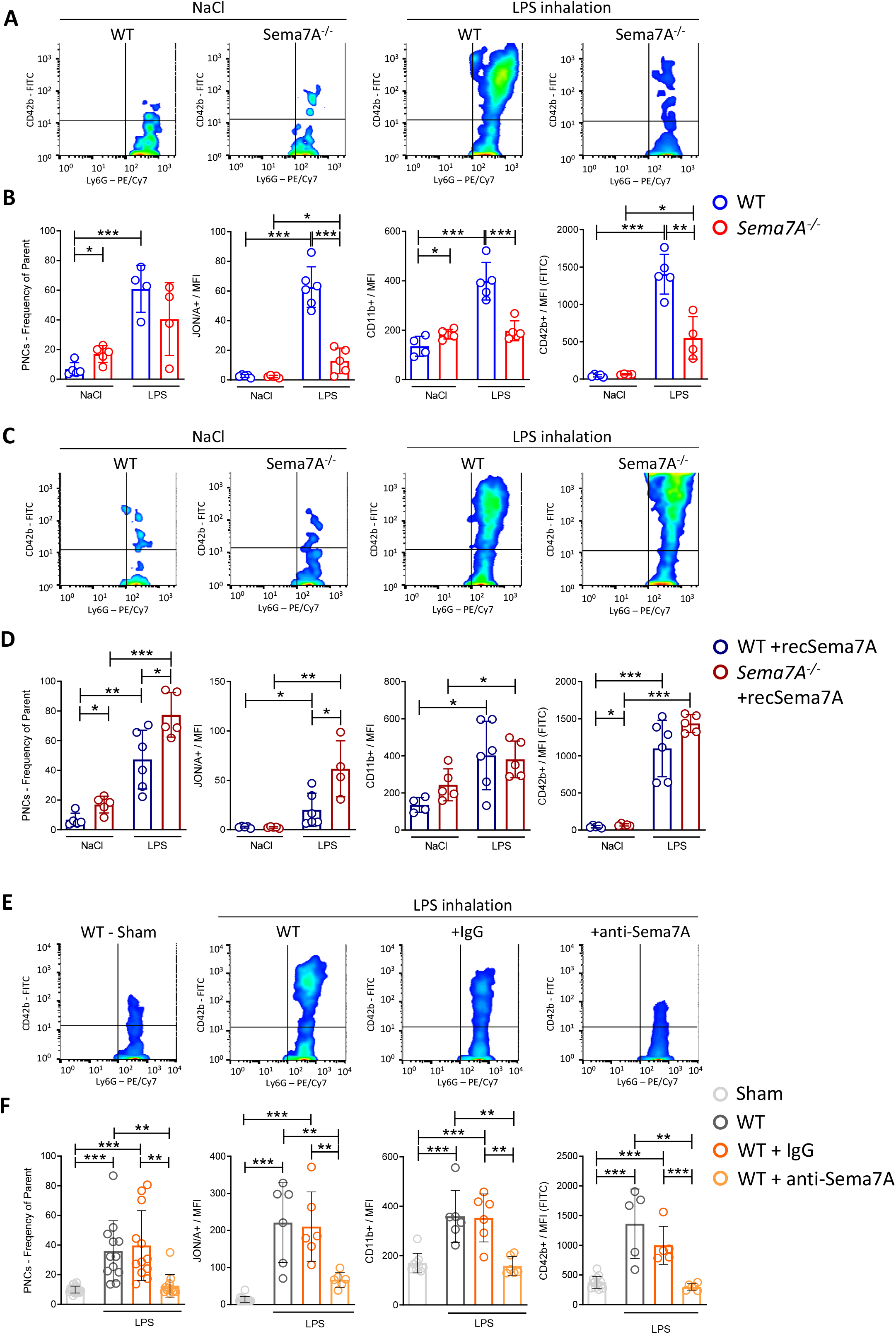
Activation of neutrophils and platelet-neutrophil complex formation is Sema7A dependent. Murine blood was collected from WT and *Sema7A*^*-/-*^ mice after LPS inhalation and analyzed by flow cytometry. **A)** Representative color dot blots of platelet-neutrophil complexes (PNCs; Ly6G^+^/CD42b^+^ events) in WT and *Sema7A*^*-/-*^ blood from NaCl (control) or LPS-inhaled mice. **B)** PNC formation, platelet effector GPIIb/IIIa expression (antibody clone JON/A MFI), PMN activity marker CD11b (MFI) expression and platelet activity marker CD42b (MFI) expression were assessed by flow cytometry in the mice described in (A). **C)** Representative dot blots of platelet-neutrophil complexes (PNCs; Ly6G^+^/CD42b^+^ events) in the blood of WT and *Sema7A*^*-/-*^ mice treated with recombinant Sema7A (recSema7A) after NaCl (control) or LPS inhalation. **D)** PNC formation, platelet effector GPIIb/IIIa expression (antibody clone JON/A MFI), PMN activity marker CD11b (MFI) expression and platelet activity marker CD42b (MFI) expression were assessed by flow cytometry in the mice described in (C). **E)** Representative dot blots of platelet-neutrophil complexes (PNCs; Ly6G^+^/CD42b^+^ events) in the blood of WT mice that were untreated or treated with IgG or the Sema7A blocking antibody (anti-Sema7A) after LPS inhalation compared to mice without conditioning (Sham). **F)** PNC formation, platelet effector GPIIb/IIIa expression (antibody clone JON/A MFI), PMN activity marker CD11b (MFI) expression and platelet activity marker CD42b (MFI) expression was assessed by flow cytometry in the mice described in (E). Group comparisons were performed by unpaired two-tailed Student’s t-tests (the data are the mean ± SD). ^*^p < 0.05, ^**^p < 0.01 and ^***^p < 0.001 as indicated.

### *Sema7A*^*-/-*^ animals show altered pulmonary defense in a model of *Klebsiella pneumoniae* infection

Bacterial invasion in the lung, the development of pneumonia and intrapulmonary inflammation are essential mechanisms during the defense against invading pathogens at the external pulmonary surface. We next examined whether Sema7A plays a role in this process. We used control and *Sema7A*^*-/-*^ animals to establish a model of *Klebsiella pneumoniae*-induced pneumonia and evaluated pulmonary inflammation 24 h after the initiation of the experiment. To precisely differentiate the effect of Sema7A on neutrophils, we assessed the sequential recruitment of neutrophils into pulmonary tissue. Then, we found that neutrophils were significantly more present in the interstitial space and significantly less present in the BALF of *Sema7A*^*-/-*^ animals than in those of littermate controls (**Figure 7A-C and I)**. We found a higher number of PNCs in the interstitial of the lungs of *Sema7A*^*-/-*^ animals **(Figure 7D)**. The inflammatory cytokines Il-1β and TNF-α were increased in *Sema7A-/-* animals, whereas IL-6 levels were unchanged **(Figure 7E-H)**. There were more colony forming units of *Klebsiella pneumoniae* present in the lung and fewer in the blood of *Sema7A*^*-/-*^ animals than control animals **(Figure 7J-M)**. When we determined the vascular permeability and the edema formed within the interstitial space and found increased edema formation in the *Sema7A*^*-/-*^animals **(Figure 7N,O)**. This increased intrapulmonary inflammation, increased alveolar-septal thickening and increased bacterial load.

**Figure 7:**
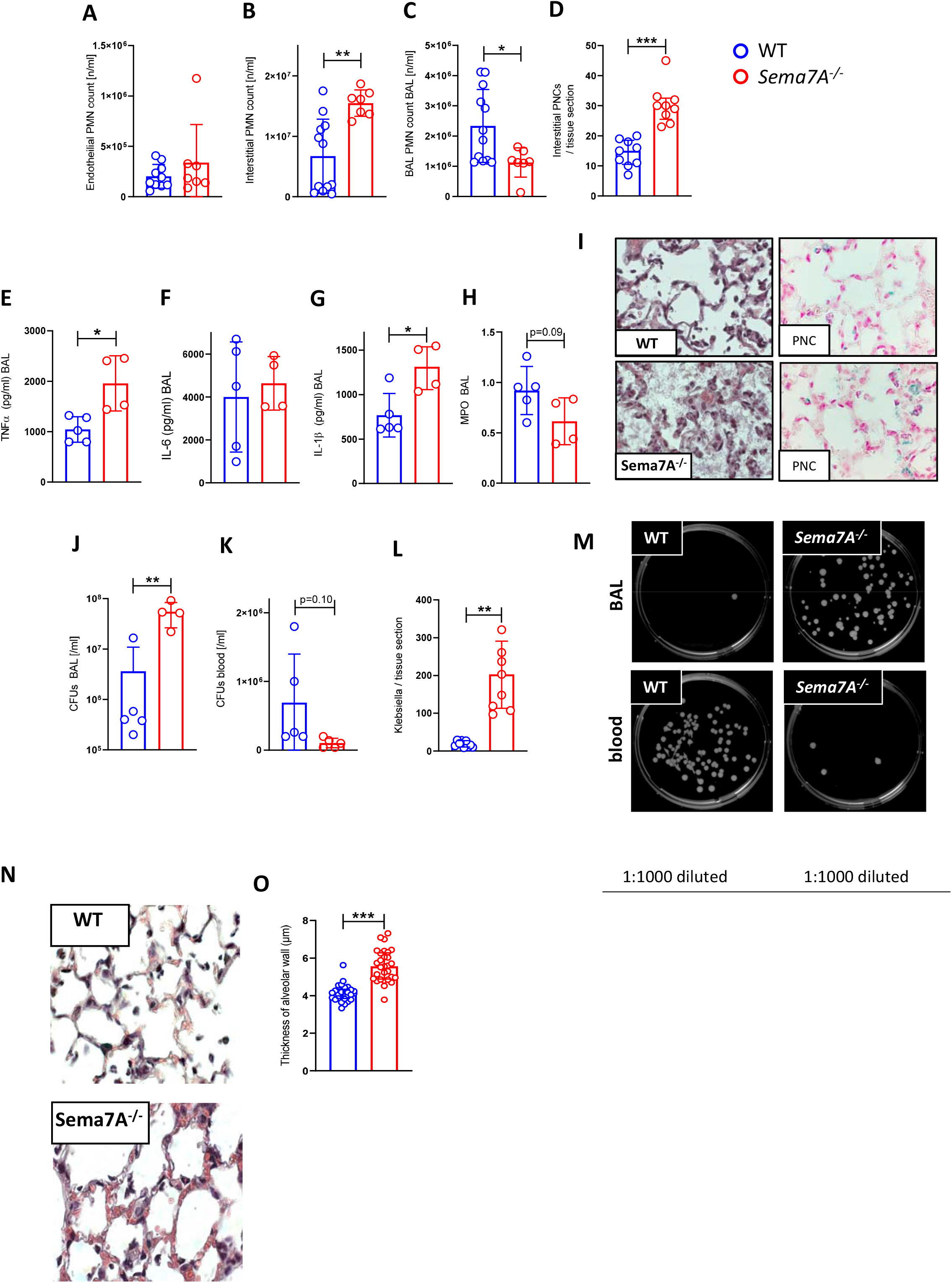
Sema7A is crucial for pulmonary defense against Klebsiella-induced pneumonia. In a murine model of bacterial-induced lung injury, 4×10^7^ gram-negative *Klebsiella pneumoniae* was administered by intratracheal instillation directly into the lungs of WT and *Sema7A*^*-/-*^ mice. Measurements of PMN counts **A)** on the endothelial surface **B)** in the interstitial space and **C)** in the BAL **D)** PNC numbers per tissue section (magnification 1000×) 24 h after Klebsiella instillation in histological lung sections of WT and *Sema7A*^*-/-*^ mice (n=3/group). The proinflammatory cytokines **E)** TNF-α **F)** IL-6 **G)** IL-1β and **H)** Myeloperoxidase activity within the BAL of WT and *Sema7A*^*-/-*^ mice **I)** Histological sections demonstrating the amount of alveolar inflammation and PNCs. Colony Forming Units in **J)** BALF and **K)** blood taken 24 h after Klebsiella instillation and incubated on nutrient agar plates for 24 h. **L)** Klebsiella counted per tissue sections and **M)** Representative images of cultured bacteria from BAL and the blood of *Klebsiella pneumoniae*-treated mice. **N)** Representative images of H&E staining and PNC immunohistochemical staining of lung tissue sections 24 h after Klebsiella instillation. (magnification 1000×; n=3/group; 3 layers were counted per mouse). **O)** Thickness of alveolar wall in tissue sections of WT and *Sema7A*^*-/-*^ mice following *Klebsiella pneumoniae* instillation (the data are the mean ± SD). ^*^p < 0.05, ^**^p < 0.01 and ^***^p < 0.001 as indicated).

**Figure 8:**
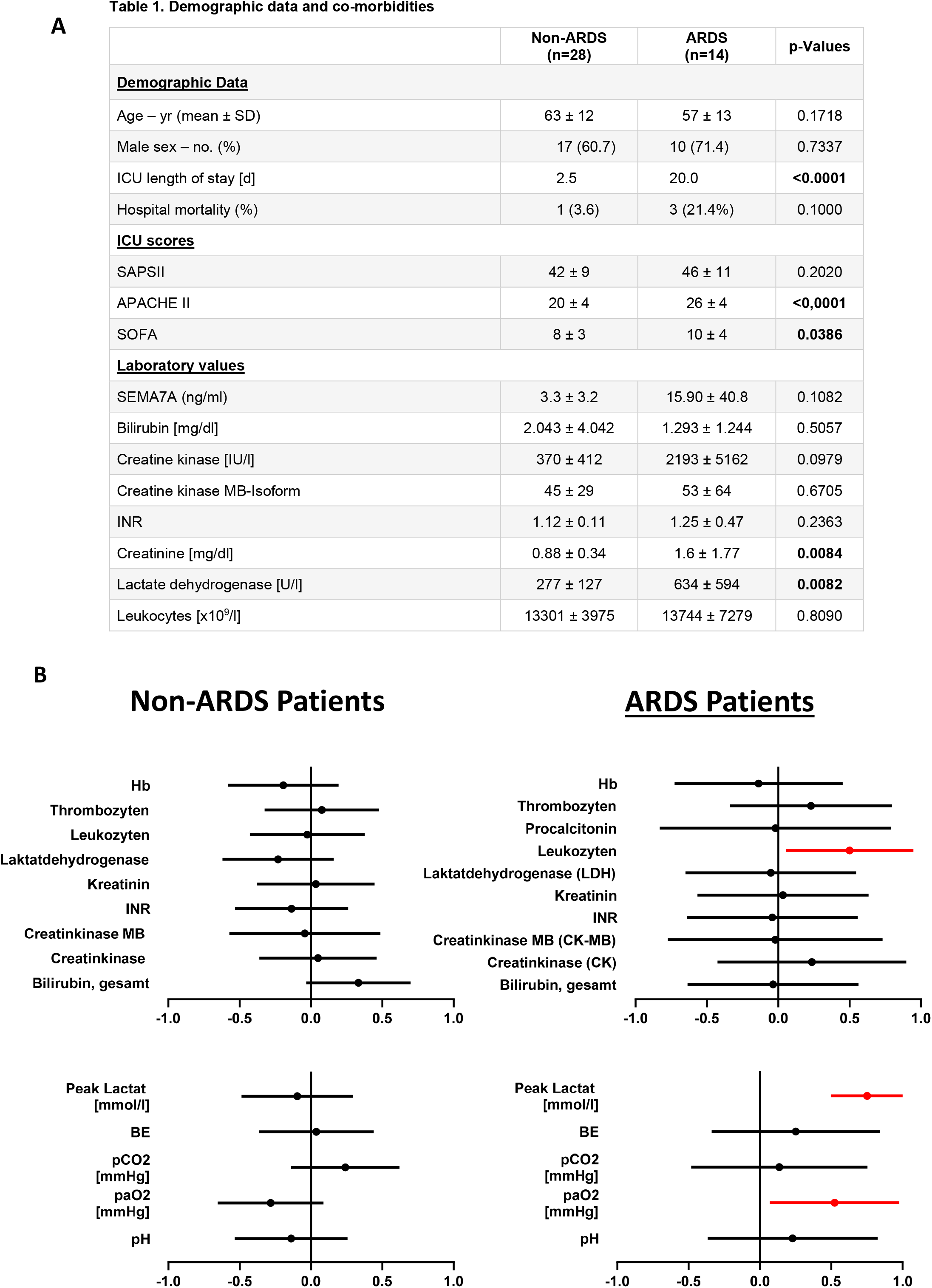
Clinical Sema7A values correlate with leukocytes and clinical oxygenation values in ARDS patients compared to ventilated controls. Demographic data and samples of patients undergoing elective surgery with postoperative ventilation and ICU stays and patients admitted to the ICU for ARDS with severe pulmonary inflammation. **A)** Demographic data, ICU scores and laboratory values of both patient groups. **B)** Correlation of different laboratory values, ventilation parameters and oxygenation values with serum levels of Sema7A.

### ARDS patients express Sema7A

In a translational approach to identifying the relevance of Sema7A in pulmonary inflammation and infection, we evaluated patients undergoing elective surgery who required mechanical ventilation and ARDS patients who had pulmonary failure due to severe pulmonary infection. We measured Sema7A blood levels on the day of admission and correlated these values with clinical and laboratory values and blood gas analysis. As expected, ARDS patients showed significantly increased values associated with illness severity, such as APACHE II and SOFA scores. Patients with ARDS showed a significant correlation between leukocyte counts and Sema7A levels. Even more interesting, the oxygenation of patients with ARDS correlated with high Sema7A levels. We did not measure neutrophil numbers in the alveolar space of these patients, since these cells are often destroyed due to the mechanical force and shear stress of ventilation, but this finding could demonstrate that patients with high Sema7A have fewer cells in the interstitial space, which affects oxygen transport through the alveolar membrane.

## DISCUSSION

Defense against external aggressors is an essential function of neutrophils that maintains the integrity and functionality of the lung. This function is important in preventing the infiltration of pathogens into the lung and eventually the human circulation. Neutrophil arrest on the endothelium and migration into the alveolar space are important early mechanisms in a multistep process. Since the treatment of pneumonia and pulmonary sepsis is not always successful, it is essential to understand the regulation or modulation of neutrophil adhesion, migration and effector functions within the lung in depth. We report here that Sema7A is an important regulator of neutrophil migration to the alveolar space and that the repression of Sema7A results in altered adhesion and delayed neutrophil migration. Sema7A activates integrins on neutrophils through the PlexinC1 receptor and influences the chemotactic behavior of these cells. In vivo, this translates into an altered immune response within the lung. As a result, Sema7A influences the early phase of inflammation, which is relevant to the outcome in murine experimental sepsis.

The activation of neutrophils is essential for the migration of these cells into the alveolar space, where they try to limit the assault on the lung from external invaders. Several proteins of the integrin class have been described to be important for this process previously described such as CD11b, PGL-1 and in addition the lectin L-selectin (*18*). The activation of L-selectin was shown to be essential for the migration of neutrophils into the lung and to limit the extent of pulmonary sepsis, the depletion of L-selectin resulted in susceptibility to pulmonary infection (*6*). Similar effects were described for CD11b and the activation of CD11b (*19*). A complex interplay between CD11b, WASP and CD42c regulates the polarization of neutrophils and attachment to microtubules on the endothelial surface (*20*). This interplay is then necessary for the meaningful and coordinated migration of neutrophils to the site of inflammation or infection. In addition, PSGL-1 is also an essential mediator of neutrophil attachment to the endothelium and is involved into the formation of platelet-neutrophil complexes (*21, 22*). This formation of PNCs also results in obstruction of the microvasculature of the lung and as a results reduces oxygenation (*23*). We have demonstrated that Sema7A alters integrin expression, increasing CD11b and L-selectin on neutrophils and reducing PSGL-1. As a result, neutrophils are activated, which essentially influences neutrophil chemotactic migration. When Sema7A is present, neutrophils are activated in a synchronistic manner and migrate across the alveolar-capillary barrier to reach the alveolar space and fight invading pathogens. When Sema7A is not present, this coordinated induction does not occur, and neutrophils migrate in an uncoordinated fashion. We observed that neutrophils in *Sema7A-/-* animals showed altered adhesion and transmigration in response to inflammatory stimuli. This alteration translated into an altered migration pattern during bacterial sepsis in the lung and neutrophil arrest in the interstitial space, where they cannot be sufficiently activated against invading bacteria. Thus, the animals showed reduced defenses and died earlier than animals with physiological Sema7A expression. PlexinC1-dependent integrin activation has not been described before in neutrophils. The role of Plexin C1 during inflammation and pulmonary inflammation was evaluated previously and showed a significant effect of Plexin C1 expression during mechanical ventilation (*24, 25*). However, the fact that Sema7A has a significant effect on CD11b, L-selectin and other adhesion integrins has not been previously shown. We were also able to demonstrate that the pathways controlling the activation of these integrins are involved in the signaling events following Sema7A binding to PLXNC1. Sema7A was shown in the past to induce cytokine storm in T-cells in a manner that was dependent on α1β1 integrin receptor in T-cells (*11*). However, we could not confirm the binding of Sema7A to β1 integrin on neutrophils and showed that Sema7A binds to the Plexin C1 receptor instead.

Previous work has demonstrated that Plexin C1 is likely involved in the migration of neutrophils and other cells, which was confirmed in models using the genetic deletion of Plexin C1 (*25, 26*). However, a mechanism underlying this finding was not evaluated in this previous work. We now show here that this occurs by the binding of Sema7A to Plexin C1. In addition, whether the soluble form of Sema7A mediates the activation of integrins, which are essential for neutrophil migration during inflammation and infection, is unclear. Recent work has demonstrated that Sema7A is essential for a coordinated sequence of events during inflammation and that Sema7A is an essential mediator of the resolution of inflammation (*27*). In accordance with this work, we also showed that Sema7A expression is important during bacterial infection. Here, we used a model of *Klebsiella pneumoniae*, while Körner et al. used a model of cecal ligation and puncture, and both showed a benefit in animals expressing Sema7A. However, one must keep in mind that the expression of Sema7A and Sema7A target receptors is organ-specific; therefore, organ-specific immune responses to inflammatory or infectious stimuli are possible in response to this protein. We have previously shown that neutrophil migration is altered through Sema7A during myocardial infarction and in hypoxic tissue inflammation (*12, 13*). During myocardial ischemia, this effect was dependent on the activation of platelet GPIb through Sema7A. During hypoxia, the induction of endothelial Sema7A resulted in increased transendothelial neutrophil migration. However, these mechanisms are not involved in the results described here, and we describe a novel mechanism of integrin activation in neutrophils through direct signaling mediated by Sema7A engaging with Plexin C1.

In summary, we have shown that Sema7A has a direct effect on neutrophil integrin signaling. This effect translates into altered adhesion and migration, which are relevant to outcomes during pulmonary infection. Data in human ARDS patients corroborate this mechanism of Sema7A-mediated altered neutrophil migration. This protein should therefore be explored further for its therapeutic potential.

## Author contributions

T.G. - performed experiments, analyzed data, wrote parts of the manuscript

D.K. – performed experiments, analyzed data, wrote parts of the manuscript

L.T. - performed experiments, analyzed data, wrote parts of the manuscript

P.B. - performed experiments, analyzed data

K.H.S. - performed experiments, analyzed data

M.K. - performed experiments, analyzed data

F.K. - performed experiments, analyzed data

K.N. - performed experiments, analyzed data

A.X. - performed experiments, analyzed data

H.H. - collected patient samples, analyzed data

T.B. - performed experiments, analyzed data

A.Z. - performed experiments, analyzed data

B.N. – designed research, analyzed data, wrote parts of the manuscript

P.R. - designed overall research plan and monitored progress, wrote the manuscript

## Supplemental Figure Legends

**Supplemental Figure 1:**
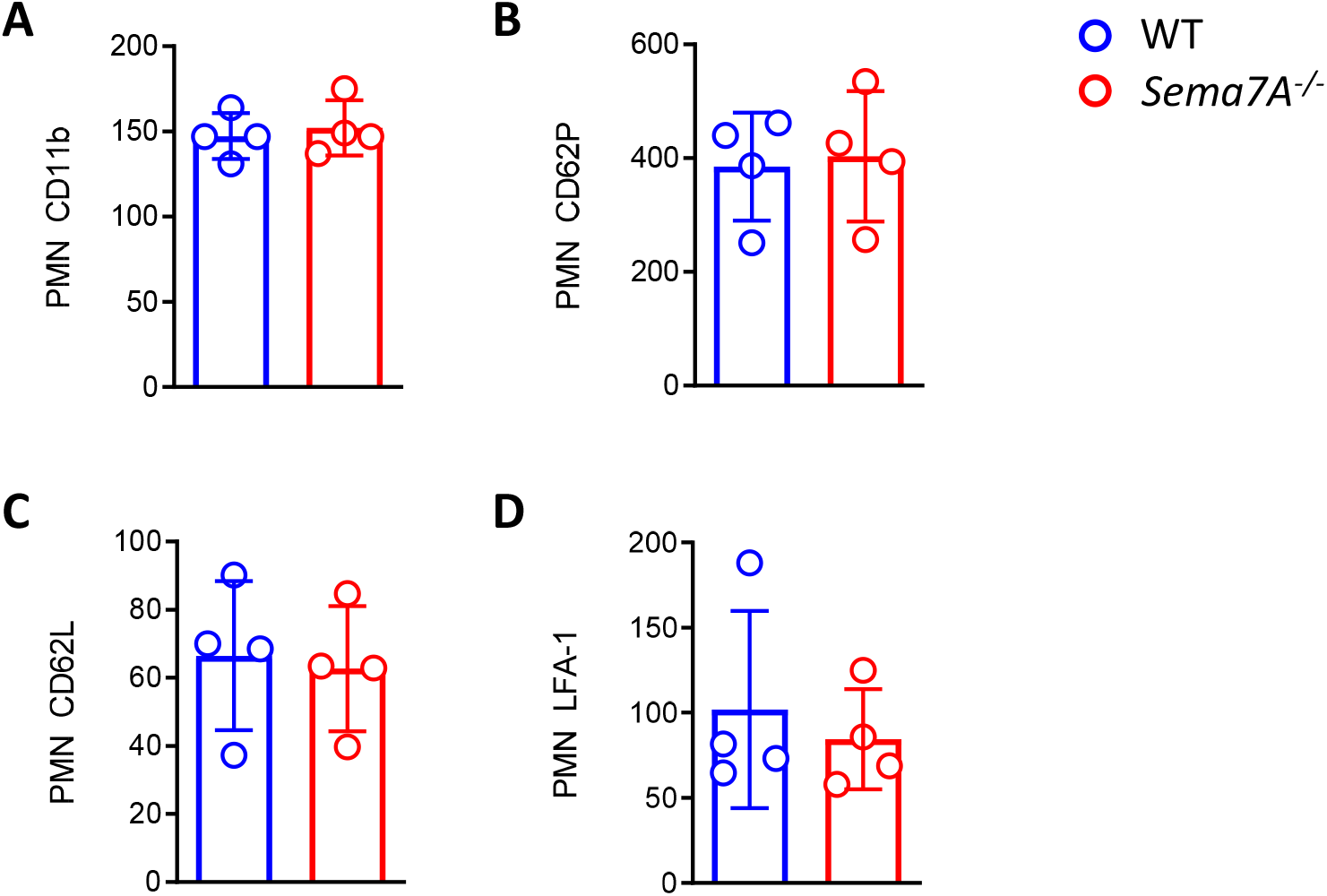
Unstimulated PMN surface expression of selected integrins. Surface expression of **A)** CD11b, **B)** CD62P, **C)** CD62L and **D)** LFA-1 on PMNs from WT and *Sema7A*^*-/-*^ animals was determined by flow cytometry. Group comparisons were performed by unpaired two-tailed Student’s t-tests (the data are the mean ± SD).

**Supplemental Figure 2:**
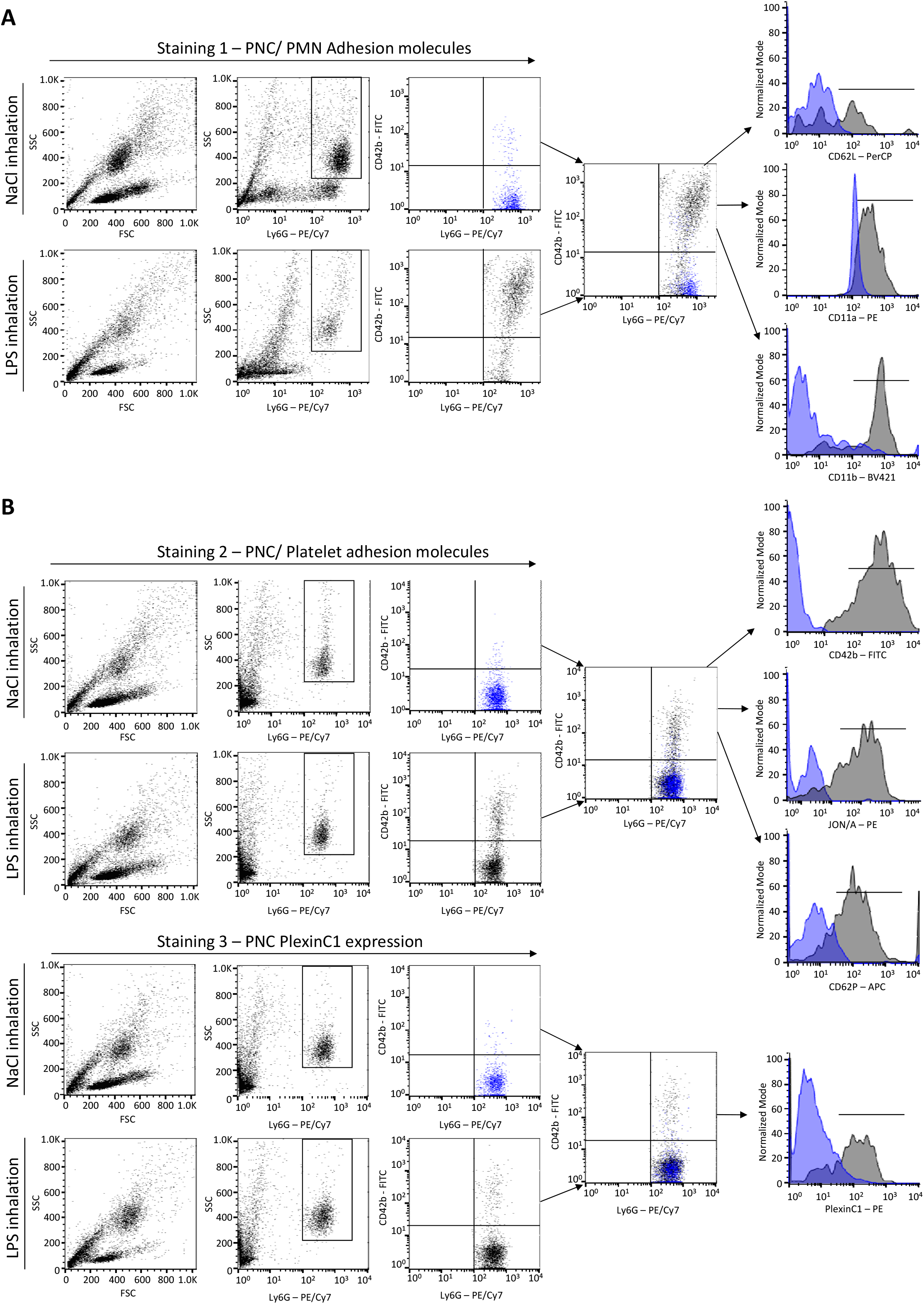
A) Gating strategy for platelet-neutrophil complexes and neutrophil evaluation *–*. Blood cell populations in NaCl- and LPS-inhaled mice were first displayed by FSC/SSC scatter plot analysis. Ly6G PE/Cy7^+^ cells (granulocytes) were gated and investigated by CD42b FITC staining to identify platelets and consequently identify platelet-neutrophil complexes (PNCs). Expression of PMN target proteins (CD62L, CD11a, CD11b) on these PNCs was determined by staining with protein-specific antibodies conjugated with PerCP, PE and BV421 (see histograms). **B) Gating strategy for platelet-neutrophil complexes and platelet evaluation *–*** Blood cell populations of NaCl- and LPS-inhaled mice were first displayed by FSC/SSC scatter plot analysis. Ly6G PE/Cy7^+^ cells (granulocytes) were gated and investigated by CD42b FITC staining to identify platelets and consequently identify platelet-neutrophil complexes (PNCs). Expression of platelet target proteins; staining (CD42b, CD41/CD61 (JON/A) and CD62P) of these PNCs was achieved with protein-specific antibodies conjugated with FITC, PE and APC (see histograms).

**Supplemental Figure 3:**
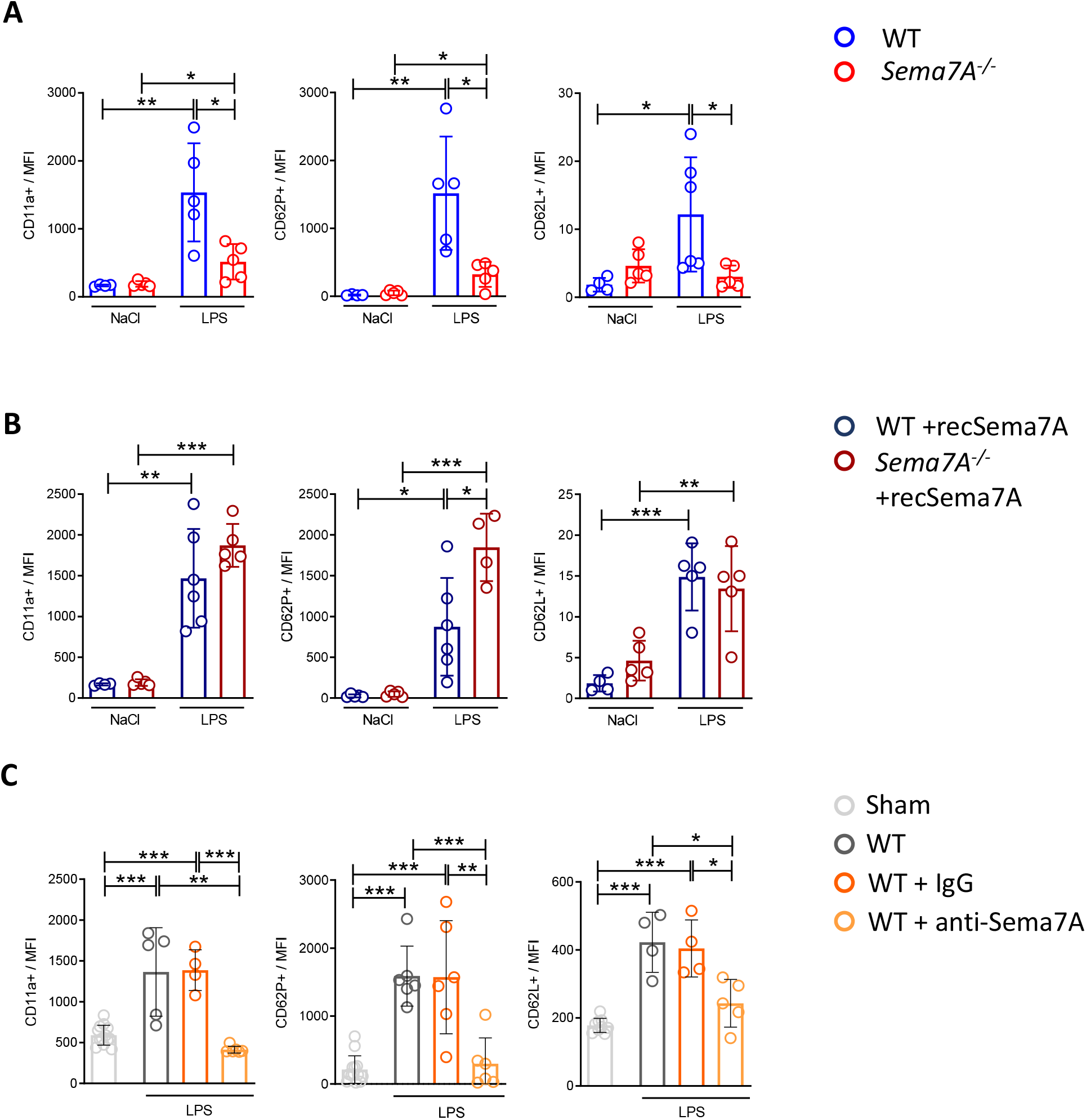
Platelet-neutrophil complex formation depends on Sema7A. Murine blood was collected from WT and *Sema7A*^*-/-*^ mice after NaCl or LPS inhalation and analyzed by flow cytometry **A)** Murine blood was collected from WT and *Sema7A*^*-/-*^ mice after inhalation of NaCl (control) or LPS (1 mg/ml) followed by a 4 h incubation time to induce lung injury and analyzed by flow cytometry. PNC surface expression of CD11a, CD62P and CD62L was measured by flow cytometry **B)** WT and *Sema7A*^*-/-*^ mice were administered recombinant Sema7A (recSema7A) i.v. and inhaled either NaCl (control) or LPS (1 mg/ml), followed by a 4 h incubation time to induce lung injury. CD11a, CD62P and CD62L surface expression on PNCs was measured by flow cytometry. **C)** WT mice were treated with Sema7A blocking antibodies or IgG i.v. and untreated WT mice were exposed to LPS followed by a 4 h incubation time. Surface expression of the target proteins CD11a, CD62P and CD62L on PNCs in the blood of these animals was evaluated by flow cytometry and compared to PNCs of untreated mice (Sham). Group comparisons were performed by unpaired two-tailed Student’s t-tests (the data are the mean ± SD). ^*^p < 0.05, ^**^p < 0.01 and ^***^p < 0.001 as indicated.

**Supplemental Figure 4:**
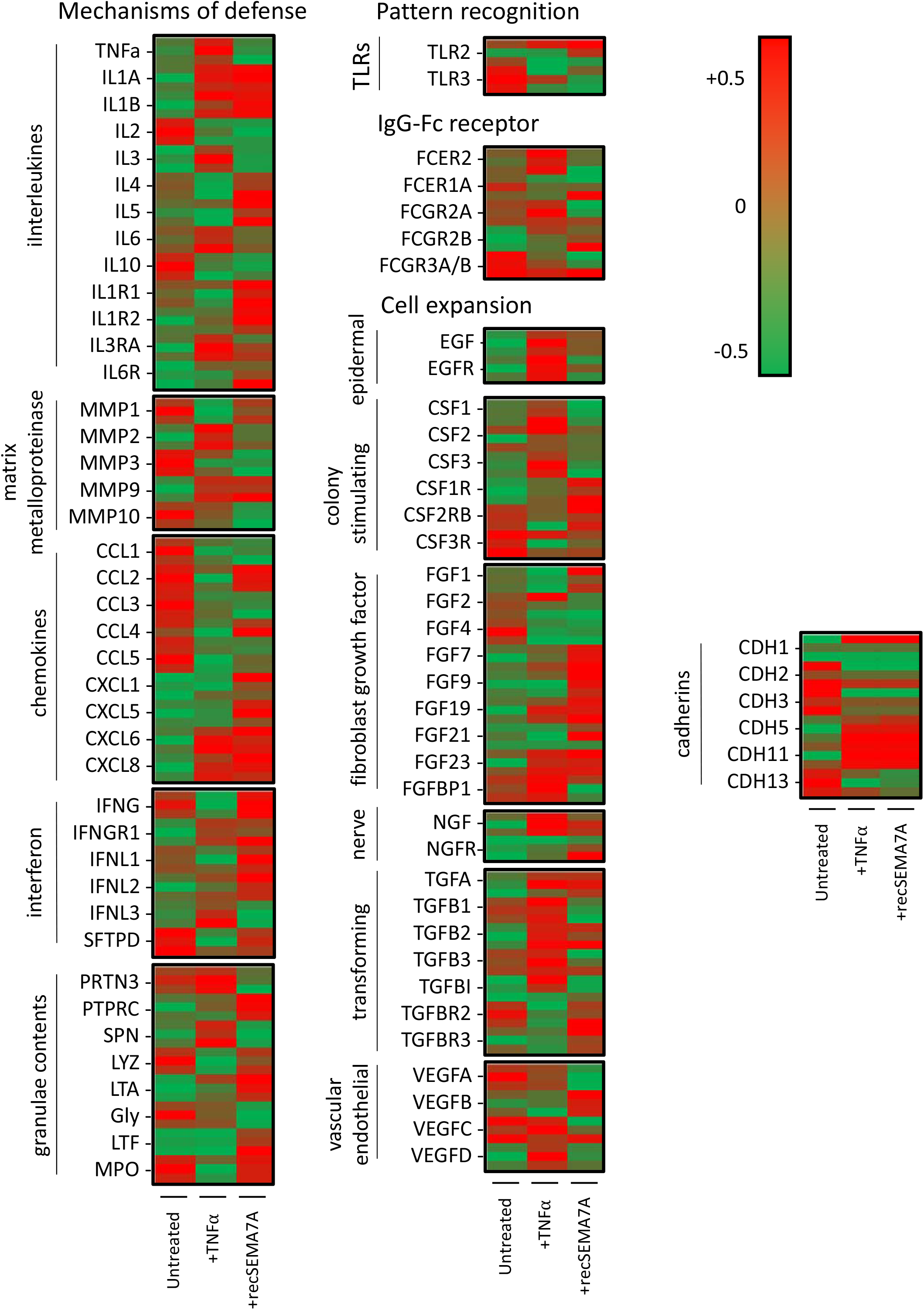
Proteomics analysis of neutrophils after stimulation with TNFα or recombinant SEMA7A. Human PMNs were incubated with NaCl, 10 ng/ml TNFα or 2 µg/ml recSEMA7A for 15 min prior to snap freezing and protein extraction for proteomics analysis. Proteins were defined as differential for |logFC| > 0.5 and an adjusted p value < 0.05. The acquired raw data were analyzed by Sciomics using the linear models for microarray data (LIMMA) package of R-Bioconductor after uploading the median signal intensities. For normalization, cyclic loess normalization was applied to all acquired data. To analyze the samples, a one-factorial linear model was fitted with LIMMA, resulting in a two-sided t-test or F-test based on moderated statistics. All presented p values were adjusted for multiple testing by controlling the false discovery rate according to Benjamini and Hochberg. Proteins were defined as differential for |logFC| > 0.5 and an adjusted p value < 0.05 from triplicate experiments. Changes in the expression of neutrophil surface proteins, membrane integrin proteins and granule proteins are displayed. Functionally, these proteins are split into the following groups: “Defense mechanisms”, “IgG-Fc receptors”, “Pattern recognition” and “Cell expansion”. The proteins in the “Defense mechanisms” group were subdivided into interleukins, matrix metalloproteinases, chemokines, interferons and granule contents, and the proteins in the “Cell expansion” group were subdivided into epidermal, colony stimulating, fibroblast growth factor, nerve, transforming, vascular endothelial and cadherins.

**Supplemental Figure 5:**
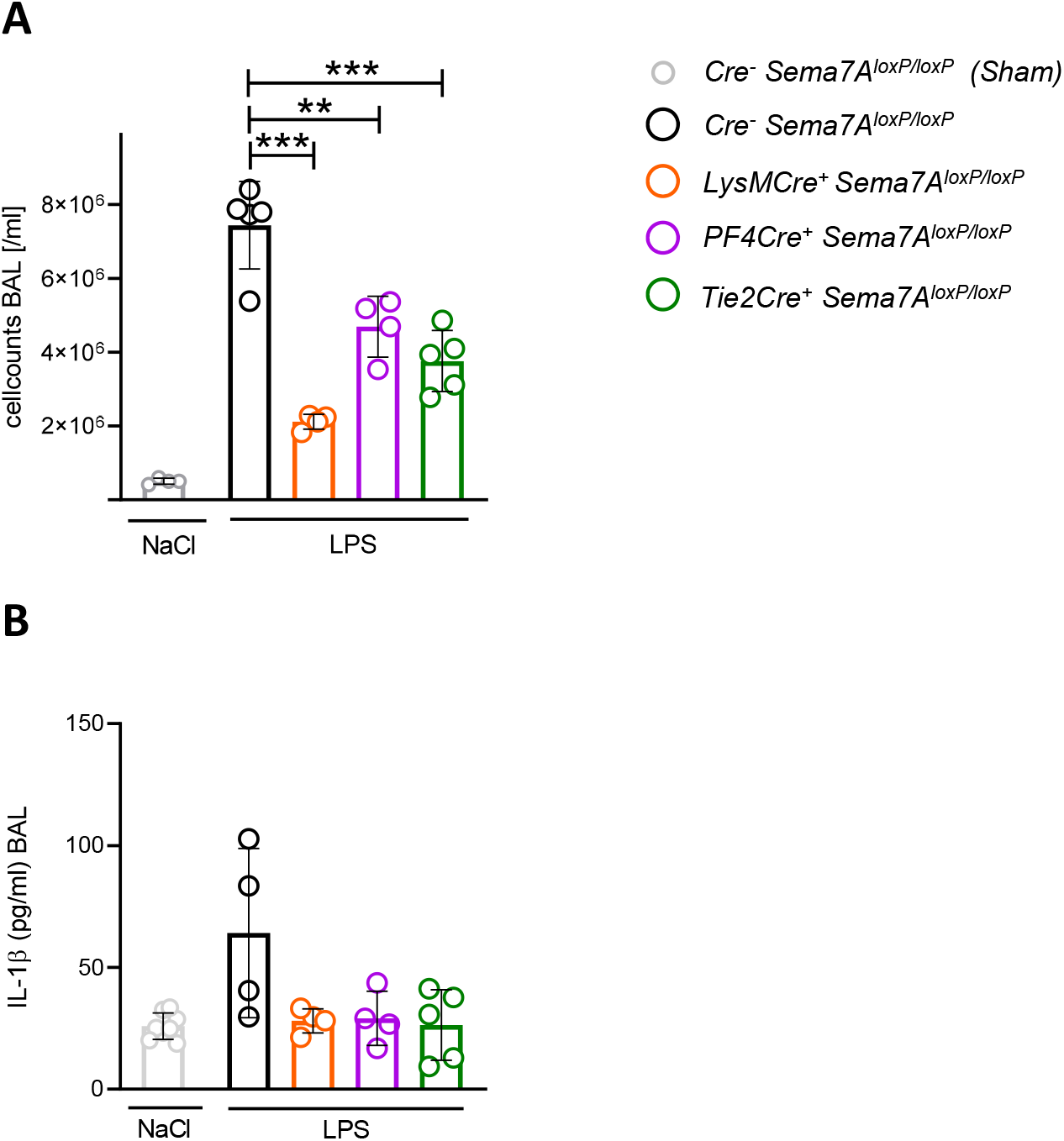
Dependence of alveolar inflammation following tissue-specific Sema7A depletion in LPS-induced lung injury. Analysis of BALF 4 h following NaCl or LPS inhalation. **A)** Cell counts in the BALF of LysMCre^+^ Sema7A^loxP/loxP^, PF4Cre^+^ Sema7A^loxP/loxP^ and Tie2Cre^+^ Sema7A^loxP/loxP^ mice compared to LPS-treated Sema7A^loxP/loxP^ Cre^-^ mice. Additionally, BALF cells from NaCl-inhaled Sema7A^loxP/loxP^ Cre^-^ Sham mice are displayed. **B)** The concentration of the cytokine IL-1β in the BALF of the (A) described mice with equal treatments. Relevant group comparisons were performed by unpaired two-tailed Student’s t-tests (the data are the mean ± SD). ^**^p < 0.01 and ^***^p < 0.001 as indicated.

**Supplemental Figure 6:**
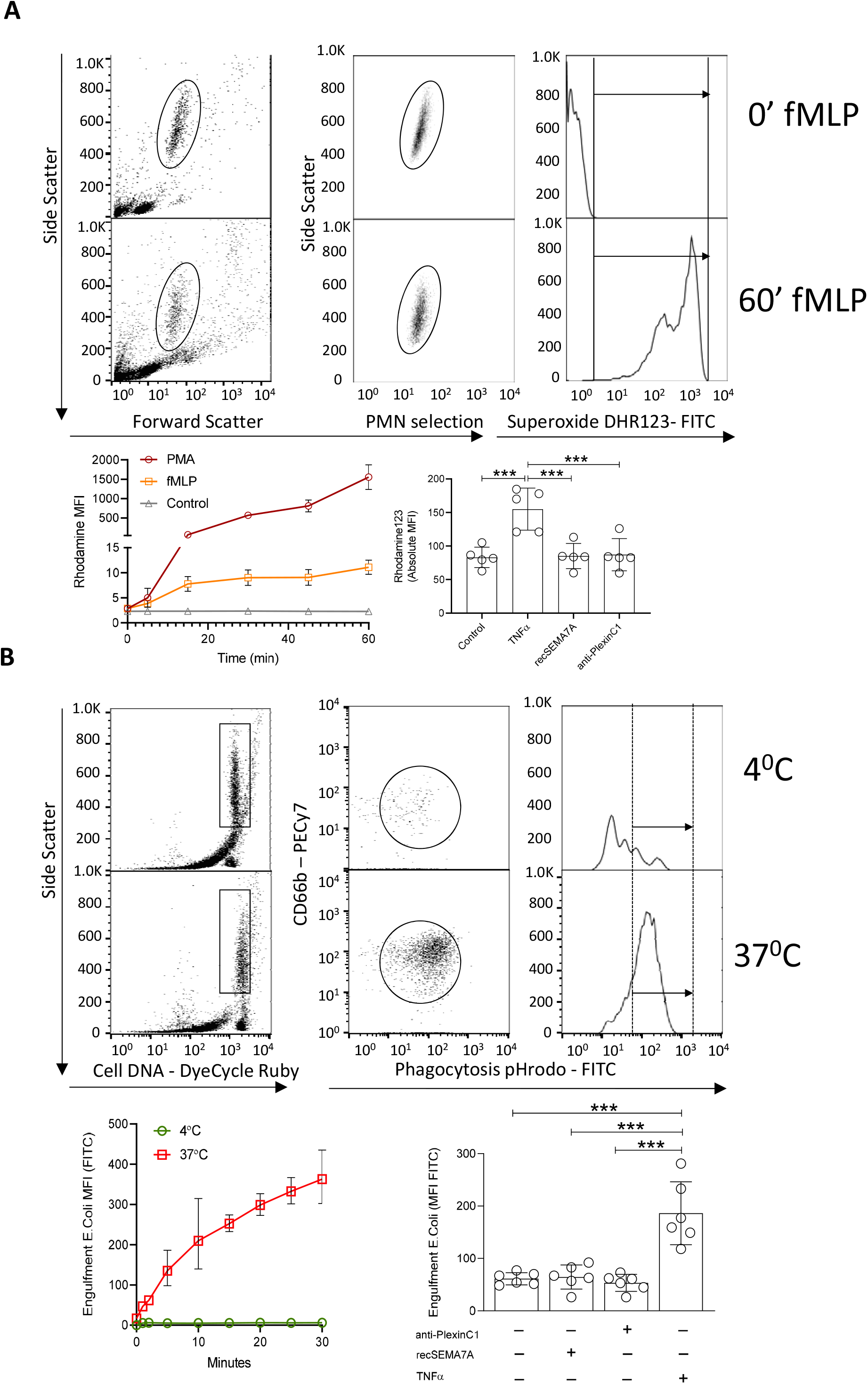
PMN superoxide production assay and PMN engulfment assay. **(A)** PMNs were identified and selected by FSC/SSC scatter, and respiratory superoxide burst was determined using an assay based on flow cytometric analysis of FITC-labeled DHR123. Pictures showing histogram examples of superoxide DHR123-FITC detection at 0 min after fMLP (control) and 60 min after fMLP treatment. As an assay control, PMNs were treated with fMLP or PMA, and a timecourse was created showing the rhodamine MFI values. Respiratory burst was not altered in PMN groups treated with recombinant Sema7A (recSema7A) or PlexinC1 antibodies (anti-PlexinC1) but was altered in the TNFα treatment group (positive control). **(B)** Isolated PMNs were first analyzed by flow cytometry to distinguish live and dead cells (Cell DNA –DyeCycle Ruby). In the FACS gating strategy used, the PMNs were marked with CD66b (PECy7), and the phagocytic capacity was determined by FITC-labeled pHrodo. A time course of bacterial uptake was performed to determine the accuracy and the optimum time for the experiment. A temperature of 4°C induced minimal phagocytosis and served as a negative control. A temperature of 37 °C was used as the optimal temperature for the experiments. The phagocytosis rate was not altered in the PMN groups treated with recombinant Sema7A (recSema7A) or PlexinC1 antibodies (anti-PlexinC1) but was altered in the TNFα treatment group (positive control). Relevant group comparisons were performed by unpaired two-tailed Student’s t-tests (the data are the mean ± SD). ^**^p < 0.01 and ^***^p < 0.001 as indicated.

**Supplemental Figure 7:**
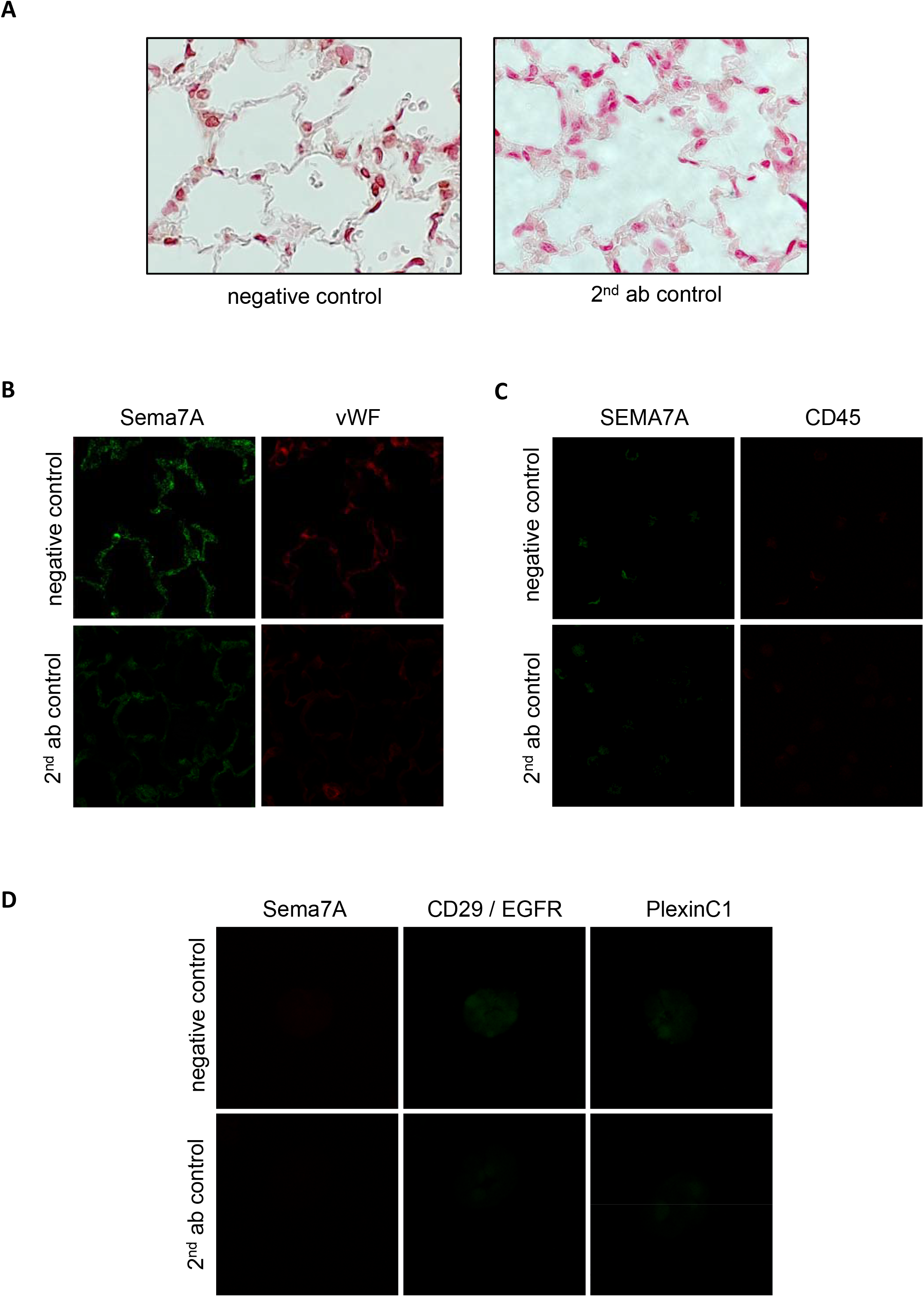
Immunohistological and immunofluorescence control staining. **A)** Immunohistochemical IgG (negative control; for Ly6G: rat; for CD41: rabbit) and secondary antibody (2^nd^ ab control) control staining of murine PNCs. **B)** Negative IgG control (for Sema7A: rabbit; for vWF: goat) and secondary antibody control for murine IF lung staining. **C)** Negative IgG control (for CD45: rabbit; for SEMA7A: mouse) and secondary antibody control for human IF PMN staining.

---

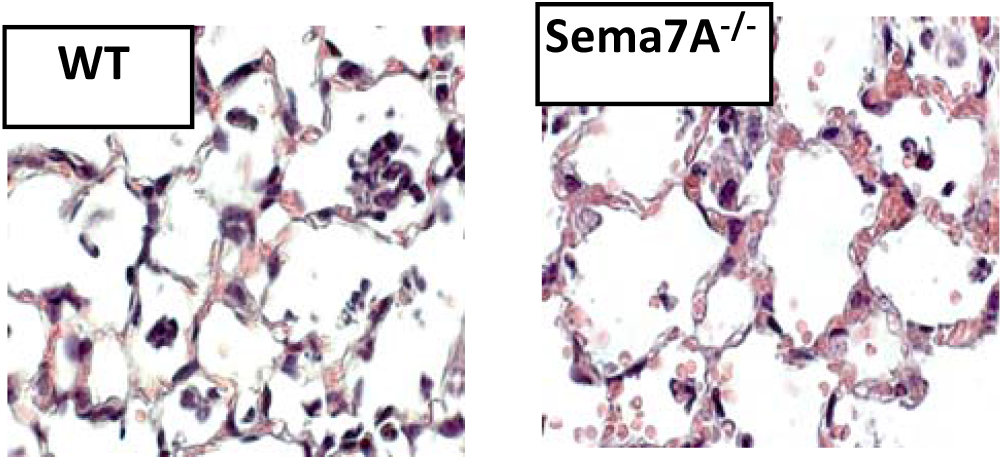

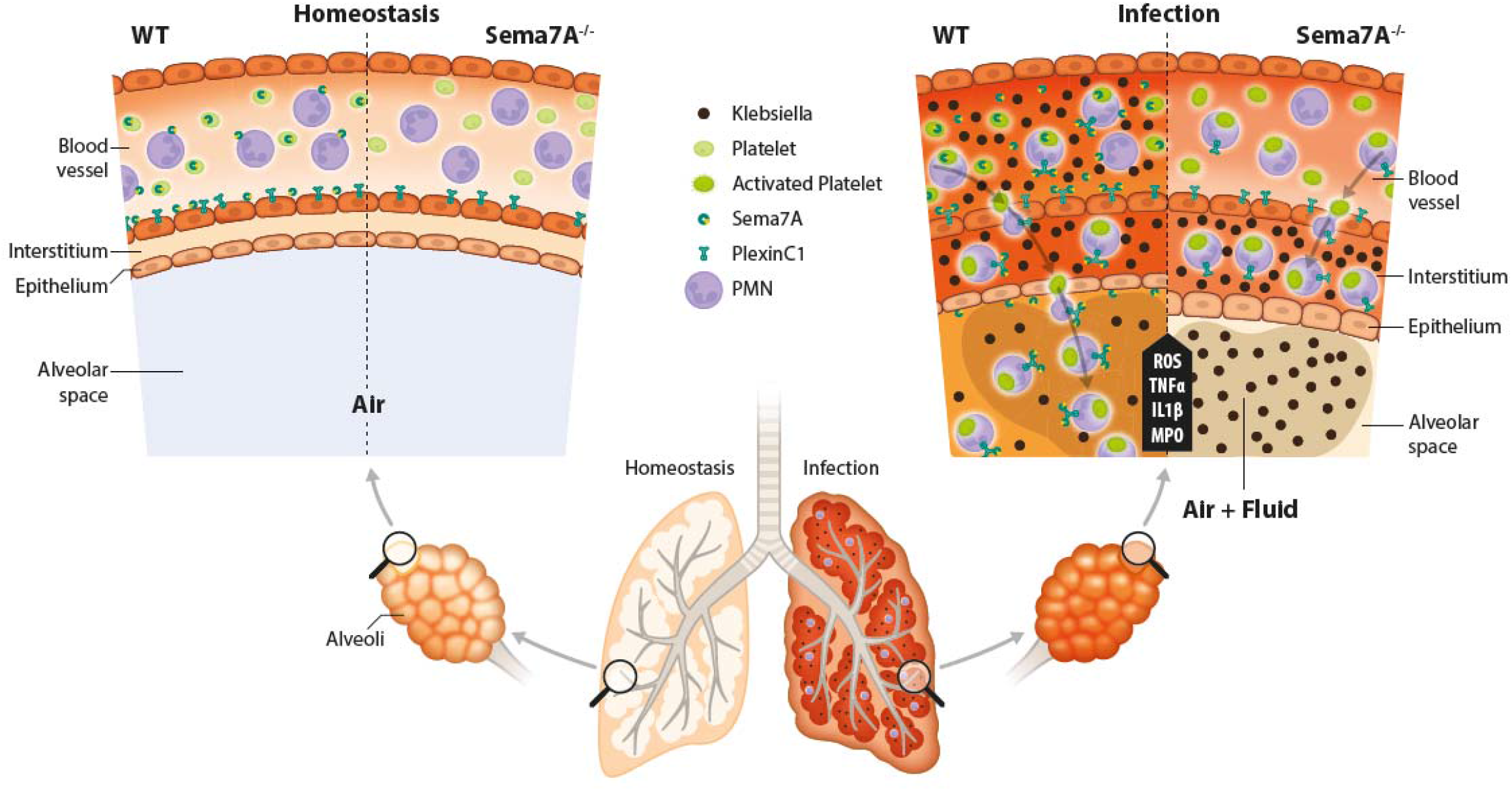

